# Hypoxic injury triggers maladaptive repair in human kidney organoids

**DOI:** 10.1101/2023.10.04.558359

**Authors:** Ana B. Nunez-Nescolarde, Yang Liao, Laura Perlaza-Jiménez, Mehran Piran, Zhengqi Cheng, Chris K. Barlow, Joel R. Steele, Deanna Deveson, Han-Chung Lee, Julie L. M. Moreau, Jinhua Li, Ralf B. Schittenhelm, Christine A Wells, Wei Shi, David J. Nikolic-Paterson, Alexander N. Combes

## Abstract

Acute kidney injury (AKI) is a common clinical disorder linked to high rates of illness and death. Ischemia is a leading cause of AKI, which can result in chronic kidney disease (CKD) through maladaptive repair marked by impaired epithelial regeneration, inflammation, and metabolic dysregulation. There are no targeted therapies for AKI or to prevent progression to CKD and insight into human disease mechanisms remains limited. Here we show that human kidney organoids recapitulate key molecular and metabolic signatures of AKI and maladaptive repair in response to hypoxic injury. Transcriptional, proteomic, and metabolomic profiling revealed tubular injury, cell death, cell cycle arrest and metabolic reprogramming in organoids exposed to hypoxia. Following return to normoxic conditions, injured organoids had increased signatures of TNF and NF-κB signalling pathways and S100A8/9, associated with maladaptive repair. Single cell RNA sequencing localized AKI and maladaptive repair markers including GDF15, MMP7, ICAM1, IL32, SPP1, C3 and CCN1 to injured tubules. Metabolic phenotypes linked to CKD were also evident, including dysregulated gluconeogenesis, altered amino acid metabolism and lipid peroxidation. iPSC-derived macrophages incorporated into organoids displayed a robust activation and inflammatory response to hypoxia. Spatial transcriptomics revealed a shift from a tissue resident-like to inflammatory macrophage states and localized effects on tubular injury and inflammation. This multi-omic analysis defines conserved mechanisms of human ischemic AKI and maladaptive repair, highlighting new opportunities to test therapeutics and model immune-mediated interactions.

## INTRODUCTION

AKI affects over 13 million people each year, resulting in high morbidity, high healthcare costs, and over 1.7 million deaths (1, 2). Ischemia is a leading cause of AKI, which can arise from cardiovascular events, urinary tract obstruction, sepsis, kidney transplantation and cardiopulmonary bypass surgery (3–5). While individuals with mild to moderate AKI can recover kidney function, they face a 5-fold increased risk of developing CKD, or of existing CKD progressing to kidney failure (6). Current management of AKI is limited to the control of blood pressure and electrolyte levels, or hemodialysis in severe cases (3, 4). Specific therapies to prevent or intervene in AKI remain an area of unmet clinical need.

During ischemia reduced blood flow limits the supply of oxygen and nutrients to the kidney causing oxidative stress and inflammation, which has a disproportionate effect on the proximal tubule (PT) (7). Injured tubules can undergo healthy (adaptive) repair via dedifferentiation and proliferation (8–10). However, sustained or severe injury can induce maladaptive repair and progression to CKD. Tubules undergoing maladaptive repair express increased levels of injury markers HAVCR1/KIM1 and GDF15, inflammatory mediators ICAM1 and CCL2, and profibrotic factors including TGFB1, HIPK2 and CCN2 which attract immune cells to the site of injury, promote fibrosis and progression to CKD (11, 12). Metabolic dysregulation is also a feature of maladaptive PT cells, marked by impaired fatty acid oxidation, reduced ATP levels and intracellular lipid accumulation (13–15). Tissue resident macrophages play an important role in AKI, honing to sites of injury, engulfing damaged cells, and activating responses that can either aid in tissue repair or promote disease progression (16, 17).

Knowledge of hypoxia-induced AKI and the transition to CKD has largely been derived from rodent models. While single cell and spatial technologies are beginning to drive insight into AKI biopsies (18), there remains a need for human experimental models to facilitate mechanistic studies and test new therapeutic strategies. The emergence of kidney organoids as a robust platform for modelling human kidney development, nephrotoxicity, cytokine stressors, and inherited disease suggests a promising avenue (19–26). Hence, we hypothesized that human kidney organoids may serve to model the tissue-intrinsic mechanisms of ischemic AKI and maladaptive repair.

## METHODS

### iPSC Culture and Organoid Generation

The following human iPSC lines were used: 522.3 (female donor; derived from GM04522 fibroblasts), 808.5 (male donor; derived from GM21808 foreskin fibroblasts). Both 522 and 808 derived by the NIGMS Human Genetic Cell Repository. D4C4 (female donor; derived from gingival tissue). Cells were maintained in Essential 8 (E8) medium (Thermo Fisher Scientific) supplemented with 1% Penicillin/Streptomycin (Pen/Strep) (Thermo Fisher Scientific) in 6-well plates coated with Matrigel (Corning, cat no. FAL35427). Cells were passaged at 70-80% confluency at 1:3 ratio using 0.5 mM EDTA (Thermo Fisher Scientific). Human iPSC were directed to differentiate into kidney organoids using the Takasato protocol (27–29), in Essential 6 (E6) medium (Thermo Fisher Scientific) with minor modifications (Supplemental Methods). Cell growth and organoid morphology was monitored by brightfield microscopy during each differentiation. iPSC cultures or batches of organoids that had slower rates of growth, unusual morphology, or poor nephron formation prior to d18 were discarded.

### iPSC-derived Macrophage differentiation

Macrophages were differentiated from PB001.1 human iPSCs as previously described (30, 31) (Supplemental Methods). Briefly, myeloid progenitors were produced from embryoid bodies after 11 days of differentiation then cultured in RPMI-1640 media (Thermo Fisher Scientific, cat. no. 11875093) with 10% FBS and 100ng/mL CSF-1 (PeproTech, cat. no. 300-25) in 6-well plates for 4 days to induce a macrophage-like state. Macrophage identity was confirmed by FACS for CD14, CD45/PTPRC and CD68 (Biolegend, cat. nos. 301830, 368512, 333816). iMacs were combined with dissociated cells from kidney monolayer differentiations at day 7 (5% iMacs, 95% kidney differentiation) and centrifuged in 1.5ml Eppendorf tubes to generate iMac-containing organoids, which were subsequently cultured as per controls.

### Hypoxic Injury

Hypoxic injury was modelled by culturing d18 organoids in 1% O_2_ for 48 h, repair was evaluated in d25 injured organoids after a 5-day recovery in 19% O_2_. Comparisons were made to stage-matched control organoids cultured in 19% O_2_. Four organoids were cultured per transwell across all experiments to minimize potential variability in the diffusion and metabolism of nutrients and oxygen.

### Bulk and Single Cell RNAseq

For bulk RNAseq, RNA was extracted from 3 replicate samples for each experimental group and sequenced using a 3’-Multiplexed method (32). Data was normalized using the counts per million (CPM) method, differentially expressed (DE) genes were defined using voom and limma R packages (33, 34) in Degust software (35) (BH adjusted p-value < 0.05 & absolute LogFC > 0.5). Enrichment analysis was performed with DAVID (36) and GSEA (37). Cell type enrichment was assessed by hypergeometric over-representation analysis comparing differentially expressed genes (FDR < 0.05) against kidney cell type gene signatures (Pod, PT, LoH, DT, Stroma, Endothelial; ∼136 genes each) derived from a developing human kidney scRNAseq reference (38), with Bonferroni correction for multiple testing. Further details on data processing, quality control and analysis are provided in Supplemental Methods.

For Single cell RNAseq, 3 iPSC-derived kidney organoids were pooled per replicate, with a total of 3 replicates per condition for d20n, d20h, d25h/n, d25n. Replicates were individually labelled with CellPlex reagents (10x Genomics) and processed on the day of collection. Gene expression and feature barcode libraries were prepared using Chromium Single Cell 5’ V3.1 chemistry (10x) and sequenced on MGITech MGISEQ2000RS hardware using MGIEasy V3 chemistry for 100 bp paired-end sequencing. Quality control and analysis was performed using the R package Seurat (v. 4.3.0) (39) and related tools, detailed in supplemental materials. All scRNAseq analysis and related code is available in a github repository https://github.com/MonashBioinformaticsPlatform/sc-hyp-org.

### Metabolomic and Proteomic Analysis

LC-MS proteomic analysis (n=4/group) was conducted on an Orbitrap Fusion Tribrid mass spectrometer (Thermo Fisher Scientific) coupled to a Dionex Ultimate 3000 RSLCnano system. Raw data were analyzed with the Fragpipe software suite (40), and LFQ-Analyst was used for downstream statistical analysis (41). Significant changes were defined in pairwise comparisons using adjusted p ≤ 0.05 (Benjamini-Hochberg method) and log2 fold change ≥ 0.5 or ≤ −0.5 thresholds. 8 replicate samples per group were profiled for untargeted metabolomic analysis to account for higher variability in this method, using a Dionex RSLC3000 UHPLC coupled to a Q-Exactive Plus MS (Thermo Fisher Scientific). Sample extraction and analysis details provided in Supplemental Methods.

### Spatial Transcriptomics

Kidney organoids +/- integrated iMacs were generated and subjected to hypoxic injury as described above. 5mm paraffin sections of samples were profiled with the 380-gene Xenium Human Immuno-Oncology panel, using the Xenium Cell Segmentation kit (10X Genomics), as per the manufacturer’s protocols. Data were imported into R, loaded into a Seurat object, and normalized with SCTransform (v. 4.3.0) (37). Principal Component Analysis was performed on the SCT-normalized data, and the top 100 components were used for UMAP visualization. Cell type identities were assigned to Xenium clusters by integrating our organoid scRNAseq reference using RCTD (42) and Seurat’s anchor-based transfer, with results cross-validated and refined by manual assessment of clusters and marker gene expression on tissue images. Pseudobulk differential expression (DE) analysis was performed on log₂-transformed, quantile-normalized, and precision-weighted data with voom (limma) (43, 44). Linear models with empirical Bayes moderation (45) were used to identify DE genes at FDR ≤ 0.05 for comparisons at the level of sample and cell lineage. Cluster-based DE was performed on the single cell level with an FDR cutoff for DE ≤ 0.2. Targeted DE analysis was performed on twenty-four manually defined Regions of Interest (ROIs) containing injured nephrons +/-iMacs in the d25h/n_iMacs organoid group, with cells selected by centroid location. The same pseudo-bulk limma-voom pipeline (FDR ≤ 0.05) was used to compare all cells, non-iMac cells, and nephron cells only between +/-iMac ROIs. Further details are provided in Supplemental Methods.

### Statistical Analysis

Details on sample number, statistical analyses and cutoff thresholds for each experiment are included in Results and Figures and/or in Methods and Supplemental Materials. All data points were included in the statistical analysis and displayed in related graphs where possible.

## RESULTS

### Hypoxia induces markers of injury and inflammation in human kidney organoids

Kidney organoids were generated from three independent human iPSC lines using established protocols (27, 29, 46) (Figure 1A). Markers of stromal, proximal and distal nephron cell types were expressed by day 18 of differentiation (d18) (Supplemental Figure 1A), which was used as a starting timepoint for subsequent experiments. d18 kidney organoids were cultured in hypoxic (1% O_2_) or ‘normoxic’ (19% O_2_) conditions for 48 hours then collected at day 20 to assess injury, or after a 5-day recovery in normoxic conditions to assess repair (Figure 1B). These experimental groups are referred to as: day 20 hypoxia (d20h), day 20 normoxic control (d20n), day 25 hypoxic injury/normoxic recovery (d25h/n or ‘injured’), and day 25 normoxic control (d25n).

**Figure 1.**
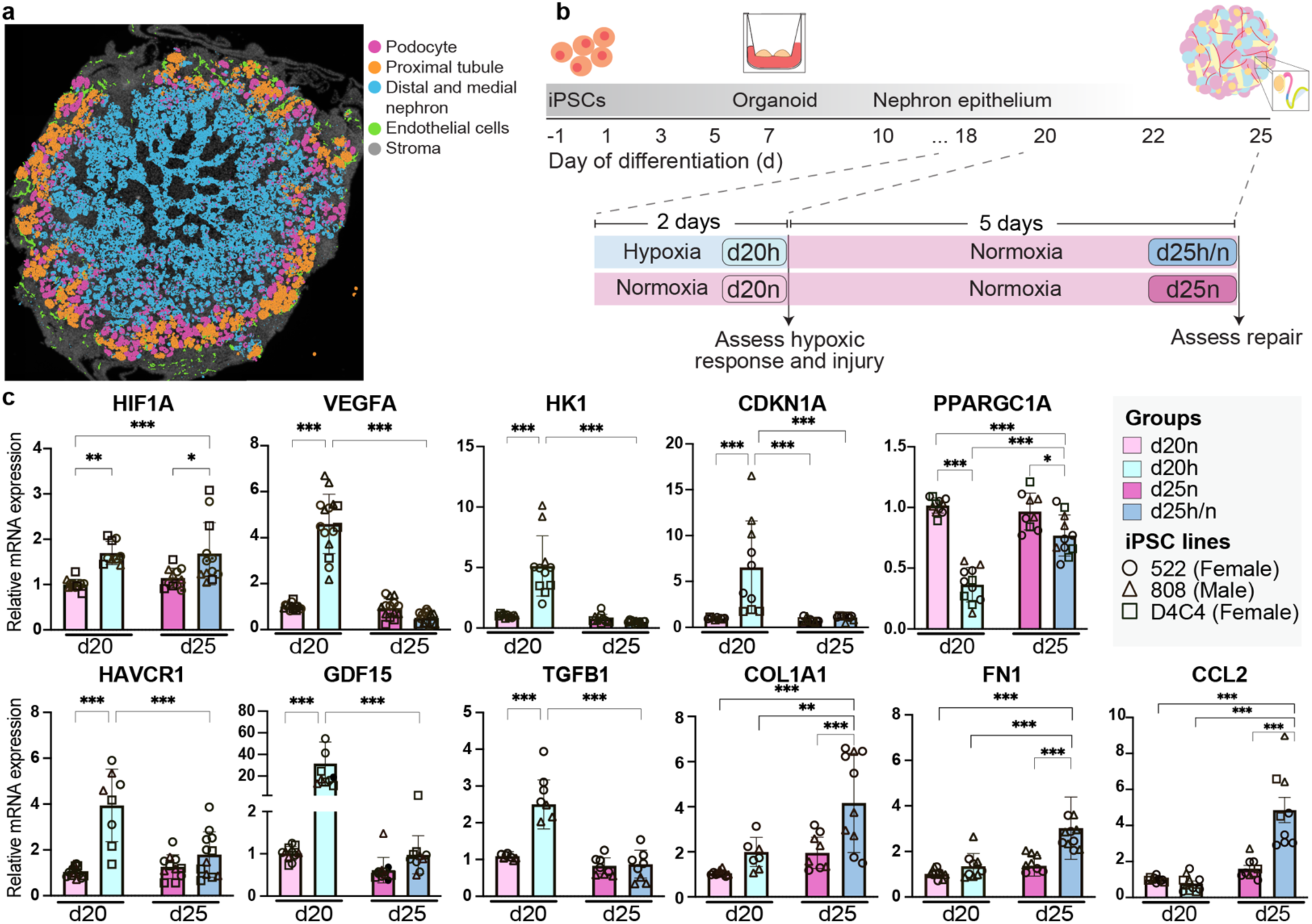
Evaluating hypoxic injury in kidney organoids from three iPSC lines. **(a)** Distribution of podocyte (pink), proximal tubule (orange), distal/medial nephron (blue), endothelial cells (green), and stroma (grey) in Xenium spatial data from a day 20 kidney organoid **(b)** Overview of experiment illustrating differentiation and organoid generation from d-1 to d18, hypoxic exposure (d18-20), normoxic recovery (d20-d25), and sample collection points at d20 and d25. **(c)** qPCR analysis of hypoxic response genes (*HIF1A, VEGFA, HK1*), markers of cell cycle arrest (*CDKN1A*), mitochondrial function (*PPARGC1A*), kidney injury (*HAVCR1/KIM1, GDF15, TGFB1*), extracellular matrix (*COL1A1, FN1*) and inflammation (*CCL2*). Data shown as mean with SD; n=7-10. Y axis shows mRNA expression normalised to *ACTB* within each sample, relative to d20n mean. Analyses were performed by one-way ANOVA with Tukey’s multiple comparison test: *p < 0.05, **p < 0.01, ***p < 0.001.

Ischemic AKI is characterized by a HIF1A-mediated hypoxic response (47, 48), tubular injury and cell cycle arrest (49). A robust hypoxic response was evident in organoids cultured in 1% O_2_ with stabilization of HIF1A protein (Supplemental Figure 1B) and increased expression of HIF1A target genes *VEGFA* and the glycolytic enzyme hexokinase 1 (*HK1*), as measured by quantitative PCR (qPCR) (Figure 1C). Mitochondrial biogenesis marker *PPARCG1A*, was significantly reduced in d20h kidney organoids, mirroring similar findings in human AKI (50). Signs of tubular damage were evident in d20h organoids with upregulation of cell cycle arrest marker *CDKN1A* (also known as p21), injury markers *HAVCR1 (KIM1)* and *GDF15*, accompanied by increased *TGFB1* which can drive profibrotic and protective responses after injury (Figure 1C) (49, 51, 52). Effective (adaptive) repair after AKI can result in a full recovery of kidney function (53–55). In contrast, maladaptive repair involves persistent expression of inflammatory cytokines such as *CCL2* and increased deposition of extracellular matrix (ECM) components including *COL1A1* and *FN1*, all of which were significantly increased in d25h/n organoids (Figure 1C).

To evaluate capacity for effective repair we reduced the duration of hypoxic exposure to 24 hours, which resulted in a hypoxic injury response (increased *VEGFA, GDF15*) without increased *COL1A1*, *FN1*, or *CCL2* expression at d25 (Supplemental Figure 1C). Thus, kidney organoids exhibit a dose-dependent response to hypoxia, effectively recovering after mild injury while prolonged exposure induces markers associated with maladaptive repair.

### Transcriptional profiling identifies markers of AKI and maladaptive repair in organoids cultured under hypoxic conditions

To gain a deeper understanding of the response to prolonged hypoxic exposure we performed bulk RNA-sequencing (RNAseq), assessing differential expression between injured and control groups on d20 and d25 (Figure 2A-C, Supplemental File 1). Top upregulated genes in d20n control organoids were associated with cell proliferation, mitochondrial metabolism, or predominantly expressed in nephron cell types in human kidney scRNAseq reference data (Figure 2B). In contrast, top upregulated genes in the d20 hypoxic group were linked to the hypoxic response, glycolysis, ER and oxidative stress (*HMOX1*), cell cycle arrest (*CDKN1A*), cell death (*JUN, FOS, BNIP3*) and inflammation (Figure 2B), with these signatures reinforced by KEGG pathway analysis (Figure 2D). Increased mitophagy, glycolysis and central carbon metabolism signatures suggest mitochondrial damage and dysfunction, alongside a shift towards oxygen-independent metabolic pathways for energy production (Figure 2D). Analysis of kidney cell type signatures among differentially expressed genes showed enrichment of distal tubule/Loop of Henle, proximal tubule, and podocyte markers in control organoids. In contrast, stromal signatures were elevated after hypoxic exposure, suggesting impaired nephron identity and stromal expansion.

**Figure 2.**
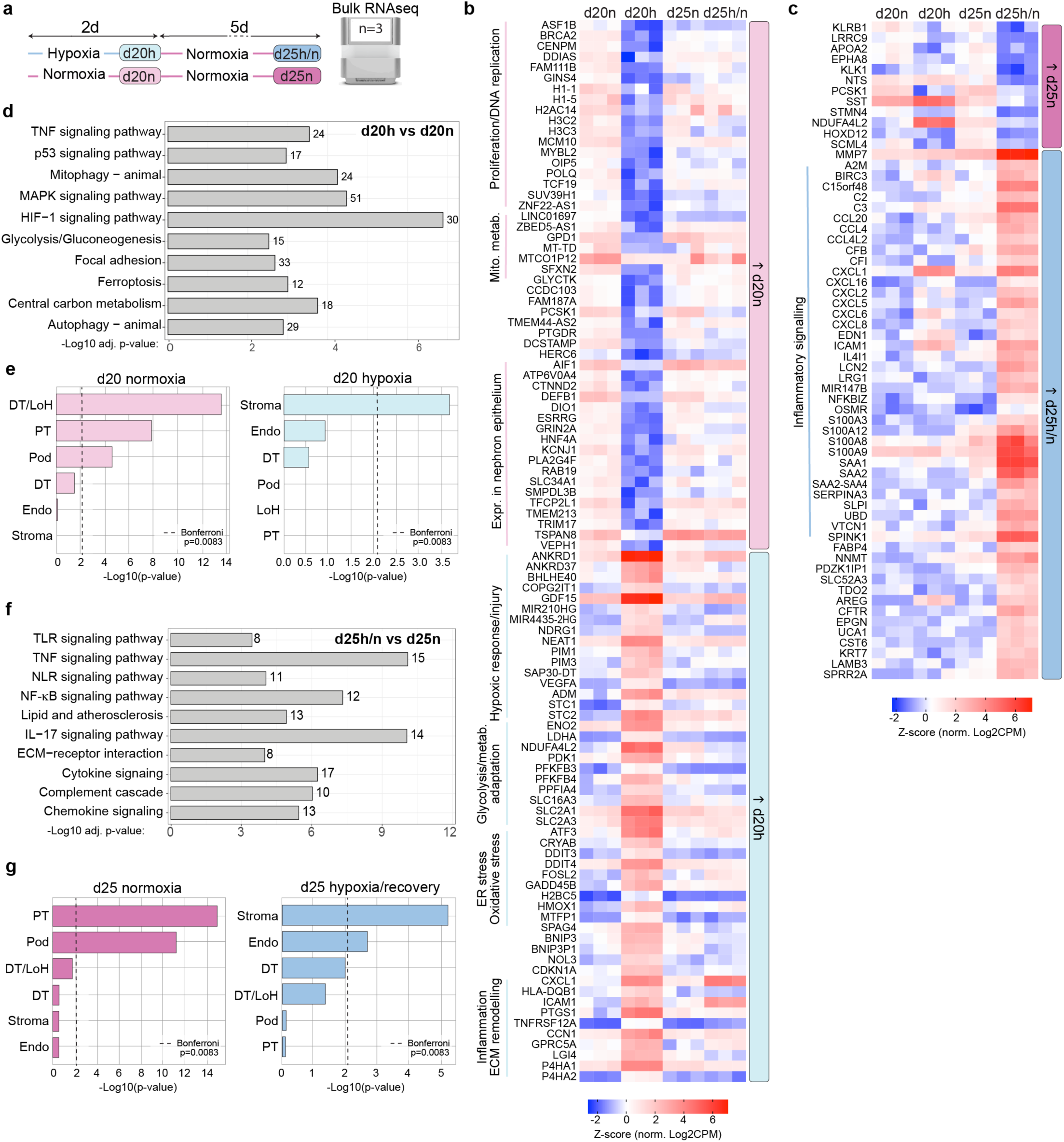
Transcriptional analysis of hypoxic injury and repair in kidney organoids. **(a)** Overview of bulk RNAseq experiment. 3 replicates were generated from a pool of 2 organoids for each group. **(b)** Top up- and downregulated genes from differential expression analysis between d20n and d20h organoids. The top 50 up and down genes (absolute LogFC>2, ranked by FDR) grouped by broad biological function or expression in scRNAseq reference. **(c)** Top up- and downregulated genes between d25n and d25h/n organoids (absolute LogFC>2, ranked by FDR), grouped by broad biological function for plotting. (**d**) KEGG enrichment analysis between d20h and d20n. Number of DE genes shown for each term. (**e**) Enrichment analysis of human kidney cell type signatures in d20 DE genes (absolute LogFC>0.58, FDR<0.05). Significance threshold (Bonferroni p<0.0083) indicated by dotted line. DT/LoH, distal tubule/Loop of Henle; PT, proximal tubule; DT, Distal tubule; Pod, Podocyte; Endo, endothelial. (**f**) Upregulated KEGG pathways in d25h/n compared to d25n organoids. **(g)** Enriched kidney cell type signatures in d25h/n and d25n organoids. Abbreviations and dotted line as per e.

Comparison of injured d25h/n organoids to stage-matched d25n controls revealed significant upregulation of genes associated with inflammatory signalling and AKI-CKD transition including *CCL2/MCP-1, CCL4/20, CXCL1/2/5/6/8/16*, *IRF1*, *STAT1/3*, *IL32*, *NFKBIA/Z,* markers of tubular injury *LCN2/NGAL*, *ICAM1, VCAM1, SPP1, LGALS3*, *S100A8/9* and extracellular matrix remodelling/fibrosis associated genes *MMP7/9, PCOLCE*, and *INHBA* (Figure 2C, Supplemental File 1). KEGG pathway analysis reinforced these findings, identifying enrichment of inflammatory programs including the TLR, TNF and NF-κB signalling pathways, interleukin and chemokine signalling, lipid-mediated inflammation, the complement cascade and extracellular matrix interactions (Figure 2F). Proximal tubule and podocyte cell type signatures were enriched in d25n control organoids, consistent with nephron maturation over time, while increased stromal and endothelial signatures in injured d25h/n organoids suggest progressive stromal expansion, early fibrotic remodelling and neovascularization (Figure 2G).

### Proteomic profiling affirms signatures of injury and maladaptive repair

To examine signatures of injury and repair at the protein level, we performed untargeted LC-MS-based proteomics on 4 replicate samples from each experimental group (Figure 3A and Supplemental Figure 2). 844 proteins were found to differ significantly across the conditions out of a total of 3630 proteins quantified (Figure 3B and Supplemental File 2). GDF15 protein was significantly increased in d20h organoids, mirroring increases observed during ischemia and tubular damage in animal models and human kidney transplants (56). GDF15 protein remained elevated in the d25h/n organoid group, reflecting reports of circulating GDF15 levels correlating with increased risk of CKD progression (57) (Figure 3C-E, Supplemental Figure 3). A hypoxic response was evident by upregulation of stress-responsive protein NDRG1 (58), the profibrotic HIF1 target CCN2 (59), and glycolytic enzymes HK1 and HK2; alongside a decrease in PCNA and phospholipid phosphatase 3 (PLPP3) (Figure 3C-D).

**Figure 3.**
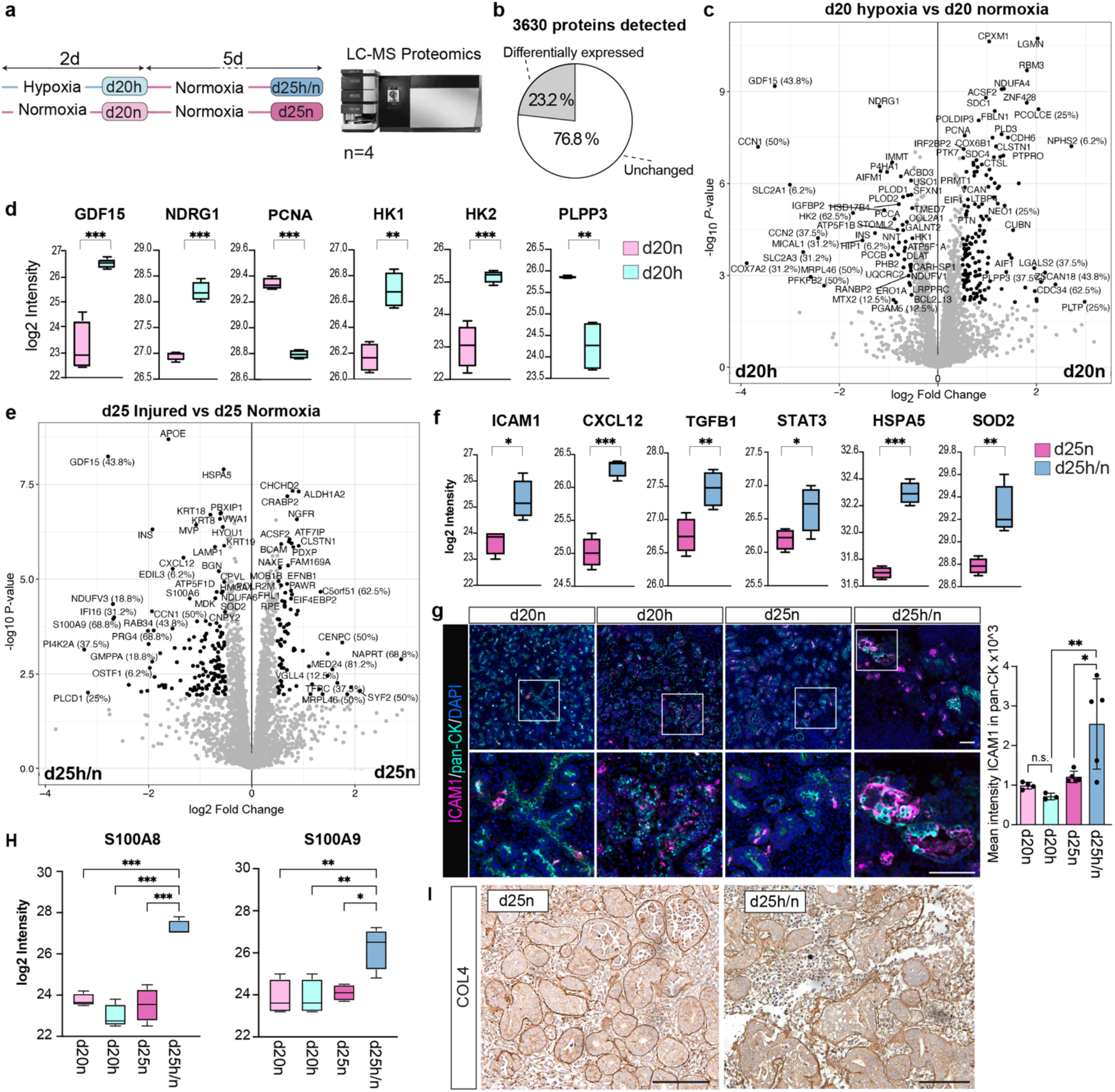
Proteomic analysis of hypoxic injury and repair. (**a**) Overview of MS-based proteomics experiment (n=4 per group). (**b**) Pie chart of differentially regulated proteins across all conditions. (**c**) Volcano plot of significantly changed proteins at d20. Percentage values indicate the extent of imputation during statistical analysis. (**d**) Intensity plots of proteins related to injury (GDF15), hypoxic response (NDRG1), cell cycle (PCNA) and metabolism (HK1, HK2, PLPP3) at d20. (**e**) Volcano plot of significantly changed proteins at d25. Percentage values indicate imputation. (**f**) Intensity plots of key proteins related to inflammatory and profibrotic processes (ICAM1, CXCL12, TGFB1, STAT3) and oxidative stress (HSPA5, SOD2) at d25. (**g**) Immunofluorescence for pan-cytokeratin (pan-CK, tubules), ICAM1 (inflammatory marker) and DAPI. Bottom panels show magnified regions indicated by white box in upper panels. Scale bars for all images shown in right hand panel = 100 μm. Quantification of mean fluorescence intensity of ICAM1 staining in tubules, n=4 replicates per experiment. (**h**) Markers of maladaptive repair S100A8 and S100A9 are upregulated in d25h/n organoids. (**i**) Representative image of collagen type 4 (COL 4) staining in d25h/n and d25n organoids. Scale bar = 200 μm. Asterisks and annotations for bar charts: n.s. non-significant, \**p* < 0.05, \*\**p* < 0.01, \*\*\**p* < 0.001.

Protein levels of inflammatory and profibrotic mediators were elevated in d25h/n injured organoids, including ICAM1, KRT7/8/18/19, APOE, S100A8/9 and CCN1 (Figures 3E-F and Supplemental Figure 4). A significant increase in ICAM1 levels and specificity to injured tubular cells was validated by immunofluorescence (Figure 3G). Proteins involved in cell stress and oxidative stress responses were also elevated - HSPA5 in d20h and d25h/n, SOD2 in d25h/n (Figure 3F), along with increased endoplasmic reticulum stress-related pathways in injured d20h and d25h/n organoids (Supplemental Figure 2D).

Tissue disorganization and nephron loss are pathological features of AKI and maladaptive repair. d25h/n organoids had a reduced number of tubular structures and disrupted tubular architecture (Figure 3I, Supplemental Figures 4,5).

Fewer differentially expressed proteins were detected by LC–MS proteomics than transcripts by RNA-seq reflecting the lower sensitivity, narrower dynamic range, and post-transcriptional regulation inherent to proteomics. Nevertheless, several changes were conserved across gene and protein profiles, revealing shared signatures of HIF pathway activation, metabolic reprogramming, and ECM remodelling during the acute hypoxic phase (d20), and increased inflammation, epithelial stress and ECM remodeling in injured organoids (d25) (Table 1). This cross-platform validation establishes high-confidence markers of hypoxic injury (GDF15, CCN1, NDRG1, SLC2A1/3, HK2, P4HA1/2; d20) and maladaptive repair (LCN2, S100A8/9, MMP7, KRT18; d25). Nephron injury is supported by a downregulation of genes expressed in podocytes and proximal tubules in human kidney scRNAseq reference data at both timepoints.

**Table 1:**
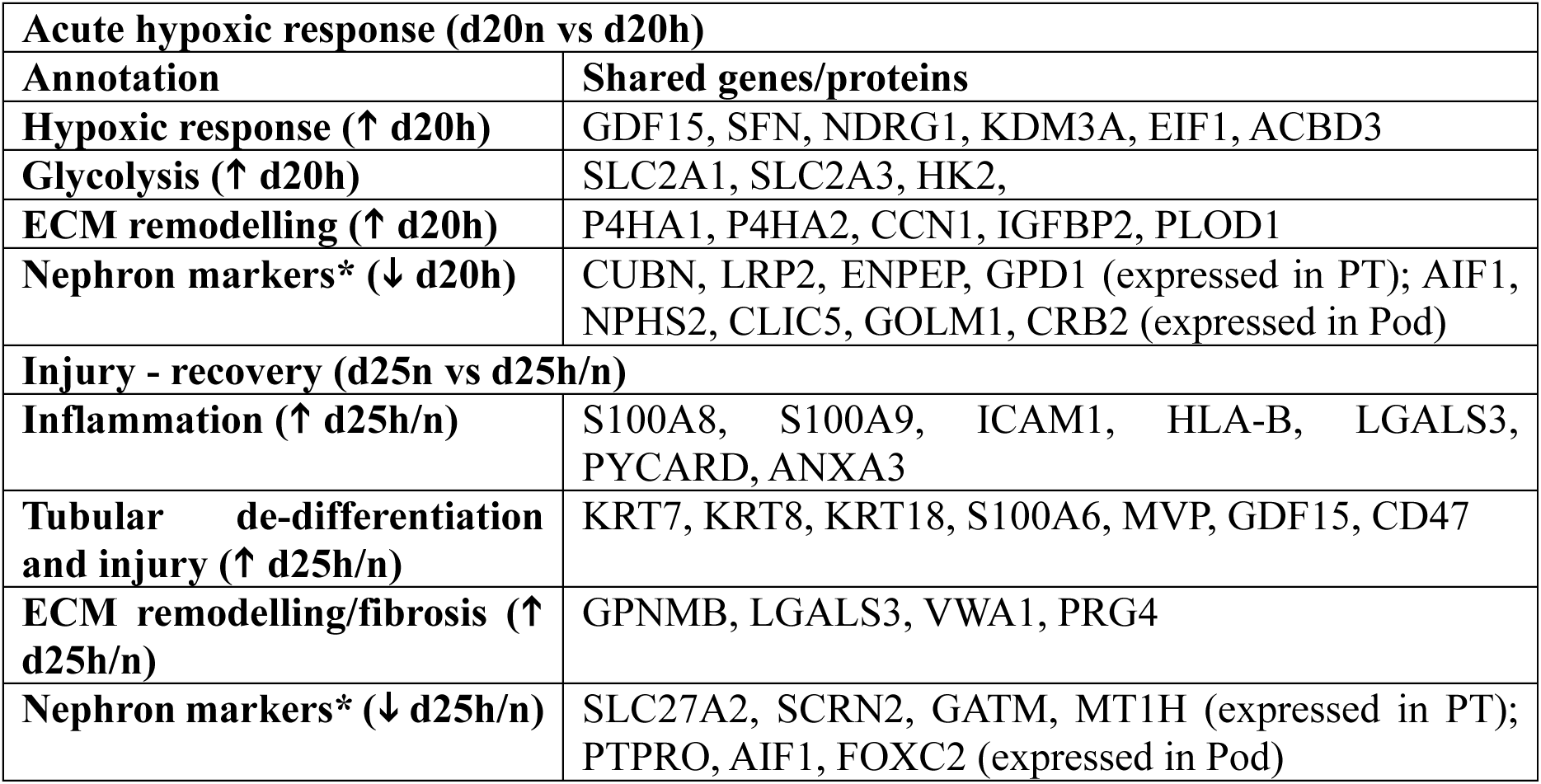
RNA-protein correlation. Overlap was calculated by identifying genes and proteins with FDR < 0.05 and absolute logFC > 0.58 in both differential datasets. *‘Nephron markers’ based on gene expression in developing human kidney scRNAseq reference data.

### scRNAseq defines changes in organoid cellular composition after hypoxic injury

We next used single cell RNA sequencing (scRNAseq) to investigate changes in cell proportions, AKI and maladaptive repair signatures. Replicate samples from each experimental group were labelled with lipid-oligo barcodes and profiled on the 10x Chromium platform (Figure 4A). Clusters reflecting expected nephron, endothelial, stromal, and off-target populations were identified by marker gene expression (38, 60) and a tool to identify human kidney cell types in scRNAseq data (61) (Figure 4B-C, Supplemental File 3).

**Figure 4.**
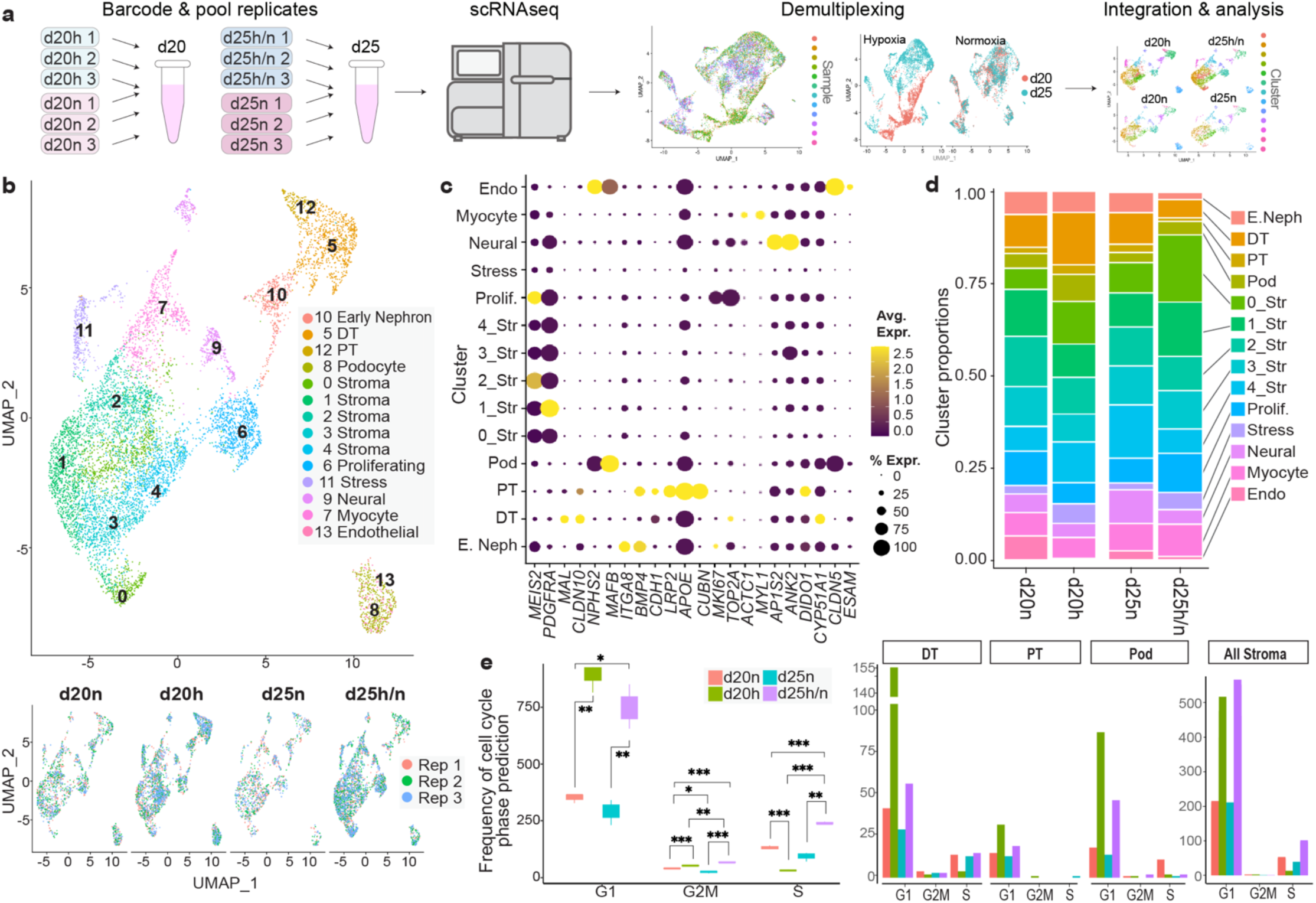
ScRNAseq analysis identifies changes in cell proportions and proliferation after hypoxic injury. (**a**) overview of scRNAseq experiment. (**b**) UMAP plot of integrated d20n, d20h, d25n and d25h/n organoid datasets coloured by cluster. Below, UMAP showing an even distribution of data from three replicates for each condition. (**c**) Dot plot of markers for each of the 14 clusters: stroma (*MEIS2, PDGFRA*), DT (*CLDN10, MAL*), podocyte (*NPHS2, MAFB*), early nephron (*ITGA8, BMP4*), PT (*LRP2, APOE, CUBN*), ‘proliferating’ (*MKI67, TOP2A*), ‘cellular stress’ (*DIDO1, CYP51A1*), muscle-like (*ACTC1, MYL1*) and neural (*AP1S2, ANK2*) and endothelial (*CLDN5, ESAM*). (**d**) Bar plot illustrating average proportions of clusters across replicates and conditions. (**e**) Box plot illustrating the frequency of cells in G1, G2/M and S phase of the cell cycle across all clusters (left), and within DT, PT, Pod and stromal clusters (right), \**p* < 0.05, \*\**p* < 0.01, \*\*\**p* < 0.001.

Cell proportion analysis (62) revealed a reduction in nephron progenitor and established nephron cell types after hypoxic injury (30% d20h to 11% d25h/n) and expansion of stromal cluster 0, which is marked by fibrosis-associated genes *SCRG1, SFRP2, KAZN* and *SCX,* (8% d25n control vs 18% d25h/n) (Figure 4D, Supplemental Tables 1-2). Analysis of cell cycle phases revealed that d20 hypoxic organoids had more cells in G1- and less in S-phase, consistent with G1 arrest, and a small but significant increase in G2/M-phase (Figure 4E). By d25, injured organoids remained G1-enriched but had an increased proportion of cells in S-phase (Figure 4E), mirroring cell cycle arrest and proliferative responses in AKI (63, 64).

### Divergent repair outcomes and shared inflammatory pathways following ischemic injury

scRNAseq analysis of hypoxic response and inflammatory signatures across all organoid groups demonstrated specificity of hypoxia, glycolysis, and kidney injury-associated genes to the hypoxic group (d20h), while inflammatory programs were enriched in the injury-recovery group (d25h/n) (Figure 5A). These signatures were increased in the nephron lineage compared to stromal or endothelial cells, reflecting the disproportionate effect of ischemic injury on the tubular epithelium (Figure 5B).

**Figure 5.**
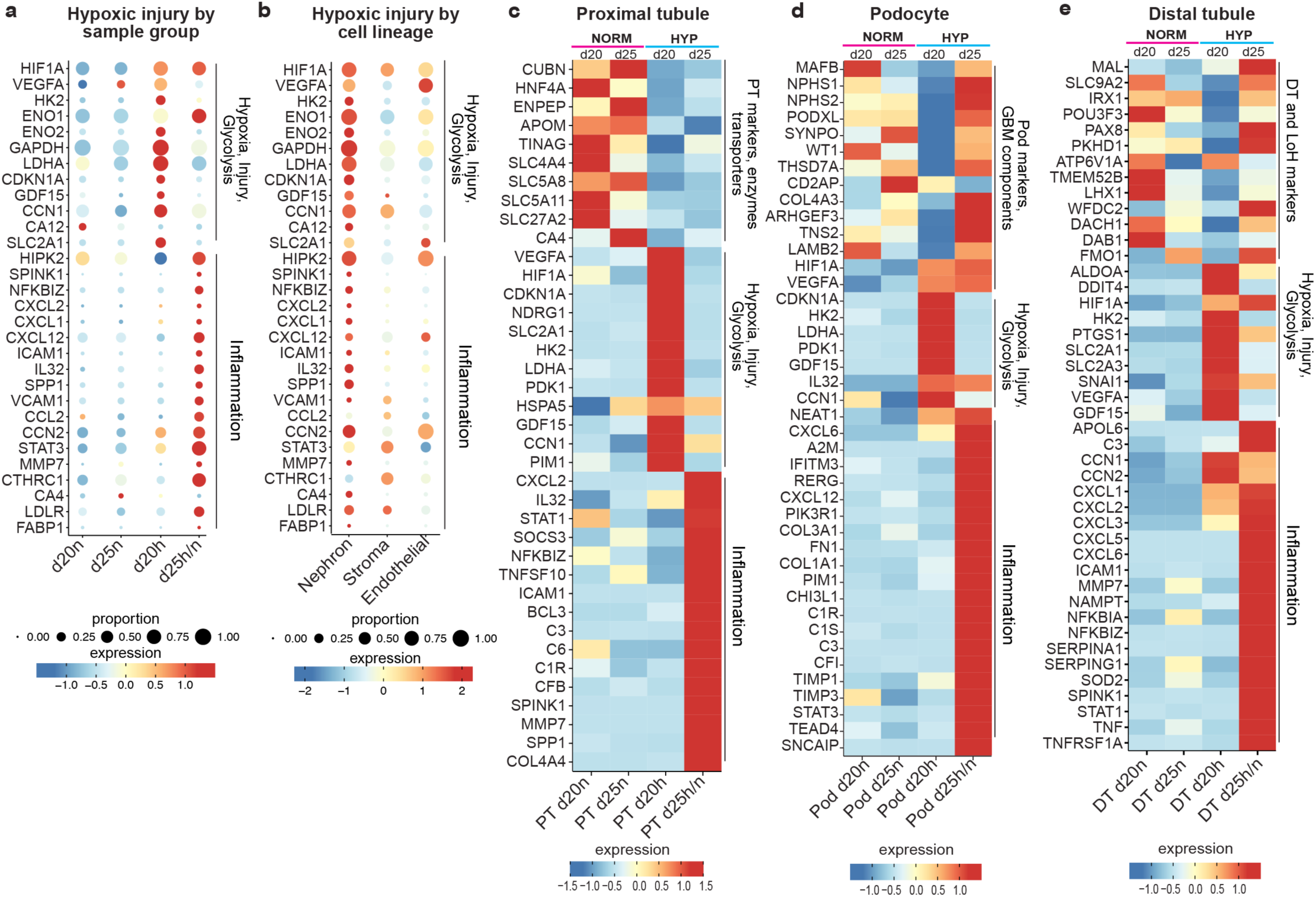
Injury and inflammation signatures are enriched in the nephron, with persistent effects in the proximal tubule. (**a**) Dot plot of hypoxia, injury, glycolysis, and inflammation–related genes across all organoid groups shows that hypoxic responses are specific to d20h, while inflammatory programs are restricted to d25h/n. (**b**) Dot plot of hypoxia, injury and inflammation signatures show enrichment in the nephron, compared to stromal and endothelial cell lineages, using expression data aggregated across all organoid groups. (**c-e**) Heat maps of selected differentially expressed genes in proximal tubule, podocyte, and distal tubule scRNA-seq clusters, including cell type markers and genes related to hypoxia, injury, glycolysis, and inflammation. Normalised expression values were scaled by each gene’s mean and standard deviation. Genes plotted have an absolute logFC > 1 and FDR < 0.05 in the d20 or d25 comparisons, except IRX1 and POU3F3, which meet an absolute logFC > 0.58 and FDR < 0.05.

We performed differential expression analysis within the proximal tubule, podocyte and distal tubule scRNAseq clusters to gain insight into injury mechanisms within nephron cell types (Supplemental File 3). Genes related to the hypoxic response, glycolysis and injury were upregulated across all clusters, accompanied by a transient reduction in cell type and functional markers. Podocyte and distal tubule markers recovered after injury, whereas several proximal tubule markers did not (Figure 5C-E). All nephron cell types retained unresolved inflammatory signatures, with common themes of complement activation, interferon and inflammatory cytokine signalling (IL32, JAK-STAT, and TNFA-NFKB), and ECM remodelling (Figure 5C-E, Supplemental Figure 6).

### Hypoxia drives metabolic dysregulation in kidney organoids

Our data suggest that hypoxic exposure triggers immediate (d20) and longer term (d25) changes in metabolism. We interrogated these changes with untargeted LC-MS metabolomics on 8 replicates per group (Figures 6A-B and Supplemental File 4). Profiles were highly conserved within groups, while substantial differences were observed between injured organoids and controls (Figure 6B).

**Figure 6.**
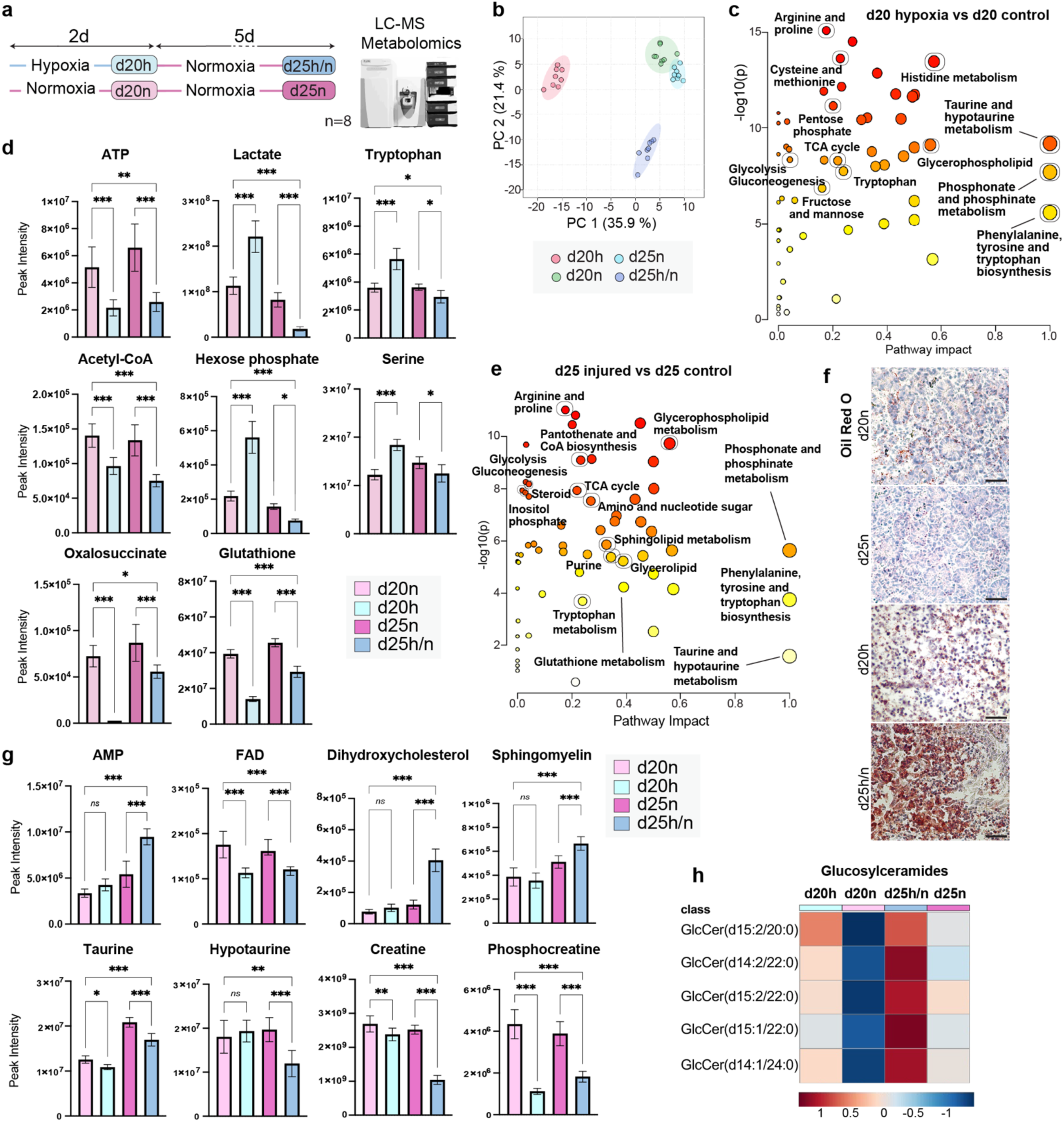
MS-based metabolomics supports metabolic dysregulation and mitochondrial disfunction during and after hypoxic injury. (**a**) Overview of metabolomic experiment with 8 replicates per condition. (**b**) PCA analysis of metabolomic data, which were normalized by the median, log10 transformed, and pareto scaled. (**c**) Metabolic pathway enrichment analysis of differences between d20h and d20n organoids. Circle size represents pathway impact score (x-axis); y-axis indicates significance, with stronger *p-*values depicted in red. Select pathways related to Ischaemic AKI are outlined and labelled. (**d**) Peak intensity of select metabolites dysregulated after hypoxia. \**p* < 0.05, \*\**p* < 0.01, \*\*\**p* < 0.001. (**e**) Pathway enrichment analysis between d25h and d25n organoids. Select pathways related to AKI-CKD transition are outlined and labelled. (**f**) Oil Red O staining of control and injured organoids at d20 and d25. Scale bars = 50 μm. (**g**) Peak intensity of select CKD-associated metabolites in d25 injured organoids. \**p* < 0.05, \*\**p* < 0.01, \*\*\**p* < 0.001. (**h**) Heatmap showing five different glucosylceramide (GlcCer) species increased in d25h/n organoids.

Pathway enrichment analysis of metabolites in d20 hypoxic and d20 control organoids indicated dysregulation of arginine and proline metabolism, cysteine and methionine metabolism, the pentose phosphate pathway, TCA cycle, tryptophan metabolism, and glycolysis/gluconeogenesis (Figure 6C). Levels of ATP were decreased while lactate levels were increased in d20h organoids, consistent with a switch to glycolysis (Figure 6D). TCA-associated metabolites acetyl-CoA and oxalosuccinate were decreased in d20h organoids (Figure 6D) consistent with reduced carbon flux into the TCA cycle during prolonged hypoxia (65). Levels of tryptophan, serine, and putative hexose-phosphate, were significantly increased in d20h organoids (Figure 6D), consistent with increased amino acid biosynthesis in hypoxia (66). Levels of the antioxidant glutathione were significantly reduced (Figure 6D).

Unbiased enrichment analysis comparing d25h/n and d25n identified disruptions in multiple metabolic pathways, including arginine and proline, glycerolipid and glycerophospholipid, sphingolipid, steroid, tryptophan, and TCA metabolism, as well as glycolysis/gluconeogenesis (Figure 6E). Consistent with these findings, Oil Red O staining revealed neutral lipid accumulation, particularly in tubular structures of injured organoids (Figure 6F). Despite 5 days of recovery in normoxic conditions, d25h/n organoids failed to restore ATP levels and instead showed increased AMP (Figure 6G), resulting in a reduced ATP/AMP ratio indicative of an energy shortage (68). Reduced levels of TCA-associated metabolites (oxalosuccinate, acetyl-CoA, and FAD) further support lasting mitochondrial dysfunction (Figures 6D, 6G).

A dyslipidaemia-like phenotype was also evident, with increased oxysterol-related metabolites, observed in oxidative stress and chronic kidney disease, and elevated sphingomyelins and ceramides associated with kidney pathology (67, 68) (Figure 6G–H). Amino acid metabolism was altered, with significant reductions in taurine, hypotaurine, tryptophan, creatine, and serine-metabolites similarly reduced in CKD patients (69–72) (Figure 6G). Collectively, hypoxia-injured human kidney organoids recapitulate metabolic features of human AKI and maladaptive repair.

### Hypoxic injury drives macrophage activation in human kidney organoids

Immune cell populations play central roles in determining kidney injury outcomes, with macrophages acting as key mediators of repair or progression to chronic disease (16, 17, 73). To begin to model immune–epithelial interactions during kidney injury, human iPSC-derived macrophages (iMacs) were combined with day 7 kidney differentiations at the point of organoid formation (5% iMac, 95% kidney), with survival of iMacs confirmed by CD68 staining on day 20 (Figure 7A-B). iMac-containing organoids and controls were then subjected to hypoxic injury (Figure 7C). Bulk RNA-seq showed no major differences in kidney cell type markers or hypoxic response genes between iMac-containing organoids and controls under normoxia or hypoxia (Figure 7D, Supplemental Figure 7, Supplemental File 5).

**Figure 7.**
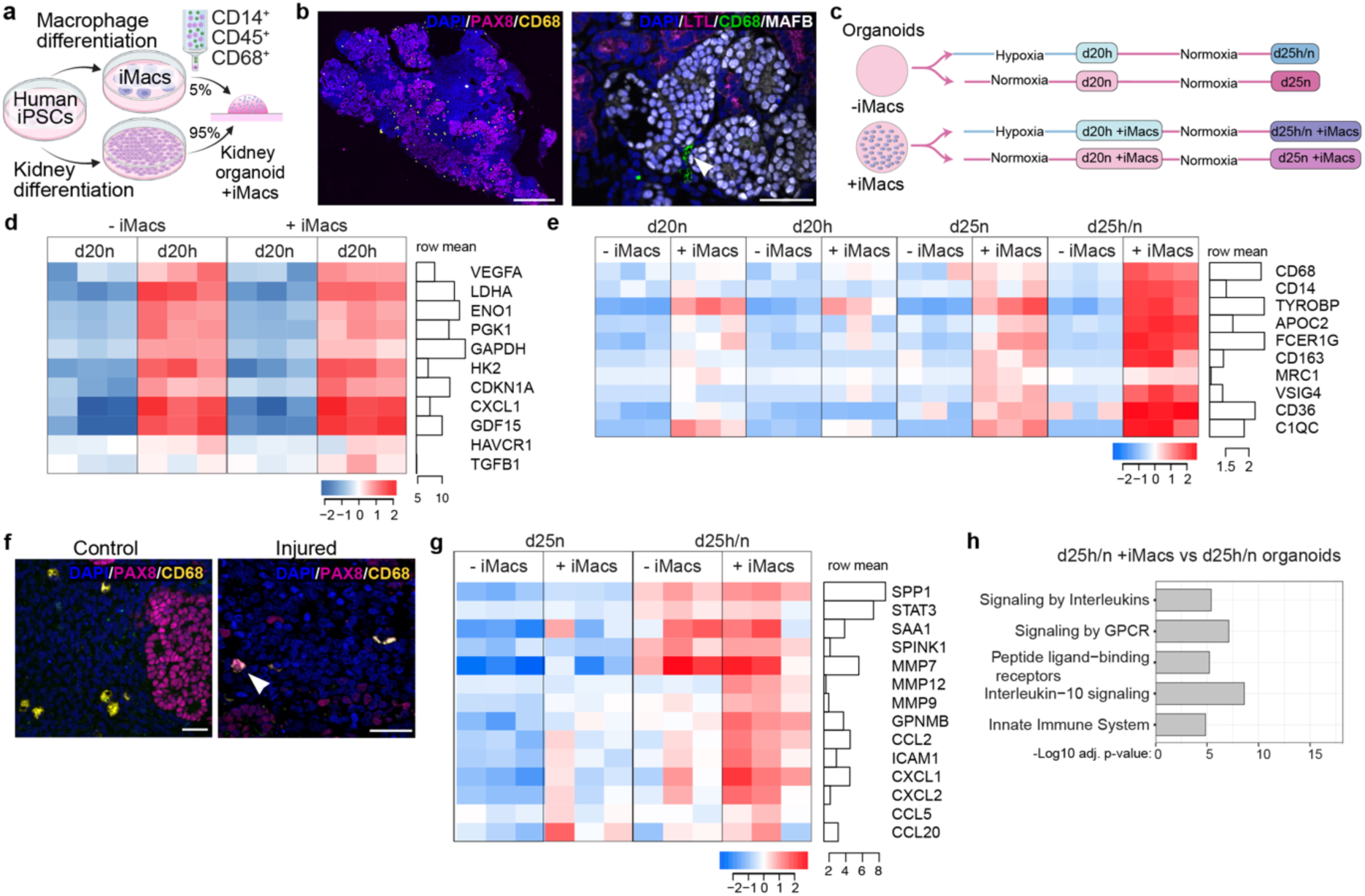
Human macrophages mount a pro-inflammatory response in injured organoids. (**a**) Overview of iMac and kidney organoid differentiation and co-culture. (**b**) Representative images of d20 kidney organoids labelled with markers of the epithelium (PAX8), proximal tubule (LTL), podocytes (MAFB), macrophages (CD68, arrowhead in right image) and nuclei (DAPI). Scale bars = 500 μm (left image) and 50 μm (right image). (**c**) Overview of hypoxic injury experiment organoids +/- macrophages exposed to 48h of hypoxia or normoxia and harvested at d20 and d25. (**d**) Heatmap of hypoxic response and injury markers in d20 organoids (Log2 CPM-normalised expression). (**e**) Heatmap of macrophage markers in d20 and d25 organoids, illustrating iMac activation in d25h/n injured organoids (Log2 CPM-normalised expression). (**f**) Representative images of d20 kidney organoids labelled with PAX8 and CD68 showing potential macrophage engulfment of a damaged epithelial cell (arrowhead); scale bars = 50 μm. (**g**) Heatmap comparing inflammatory markers in d25 organoids (Log2 CPM-normalised expression). (**h**) Upregulated pathways in d25h/n iMacs-containing injured organoids compared to injured controls (FDR < 0.05).

Following hypoxic injury, d25h/n iMac-containing organoids showed significant upregulation of macrophage activation markers (Figure 7E) without an apparent change in macrophage numbers (Supplemental Figure 7). Immunofluorescence revealed PAX8 internalization in CD68⁺ iMacs, consistent with phagocytosis of injured epithelial cells (Figure 7F). Injury-associated epithelial markers (*SPP1, STAT3, SAA1, SPINK1, MMP7*) were elevated in injured organoids whether they contained iMacs or not. However, iMac-containing organoids expressed significantly higher levels of proteins linked to tissue remodeling (*MMP9, MMP12*, *GPNMB)*. Pro-inflammatory cytokines and chemokines (*CCL2, CCL5, CCL20, CXCL1/2*) and *ICAM1* were also upregulated, along with enrichment of innate immune and interleukin signaling pathways (Figure 7G,H). Together, these data indicate that iMacs mount a pro-inflammatory response in injured organoids, recapitulating key features of macrophage activation during AKI.

### Spatial transcriptomic analysis of macrophage influence on injury and repair

To further interrogate how macrophages respond to hypoxia and affect the injury response at a cellular level, we profiled injured iMac-containing organoids and controls using the Xenium Human Immuno-Oncology (I/O) panel. The combined dataset contained 835,557 cells from 25 samples and 6 experimental groups (d20n, n=2; d20h, n=3; d25n, n=6; d25h/n, n=5; d25n+iMacs, n=4; d25h/n+iMacs, n=4; Figure 8A, Supplemental Figure 8). Clustering and scRNAseq-guided annotation of the Xenium data defined expected nephron, stromal, endothelial and macrophage cell types (Figure 8B,C, Supplemental File 6). Nephron clusters with distal and connecting segment signatures were identified (C02, C13, C20, C21), alongside proximal tubule (C11), podocyte (C04) and injured-nephron epithelium (C12), marked by *EPCAM*, *TGFB1, CDKN1A*, *ICAM1*, and *ACTA2* (Figure 8C).

**Figure 8.**
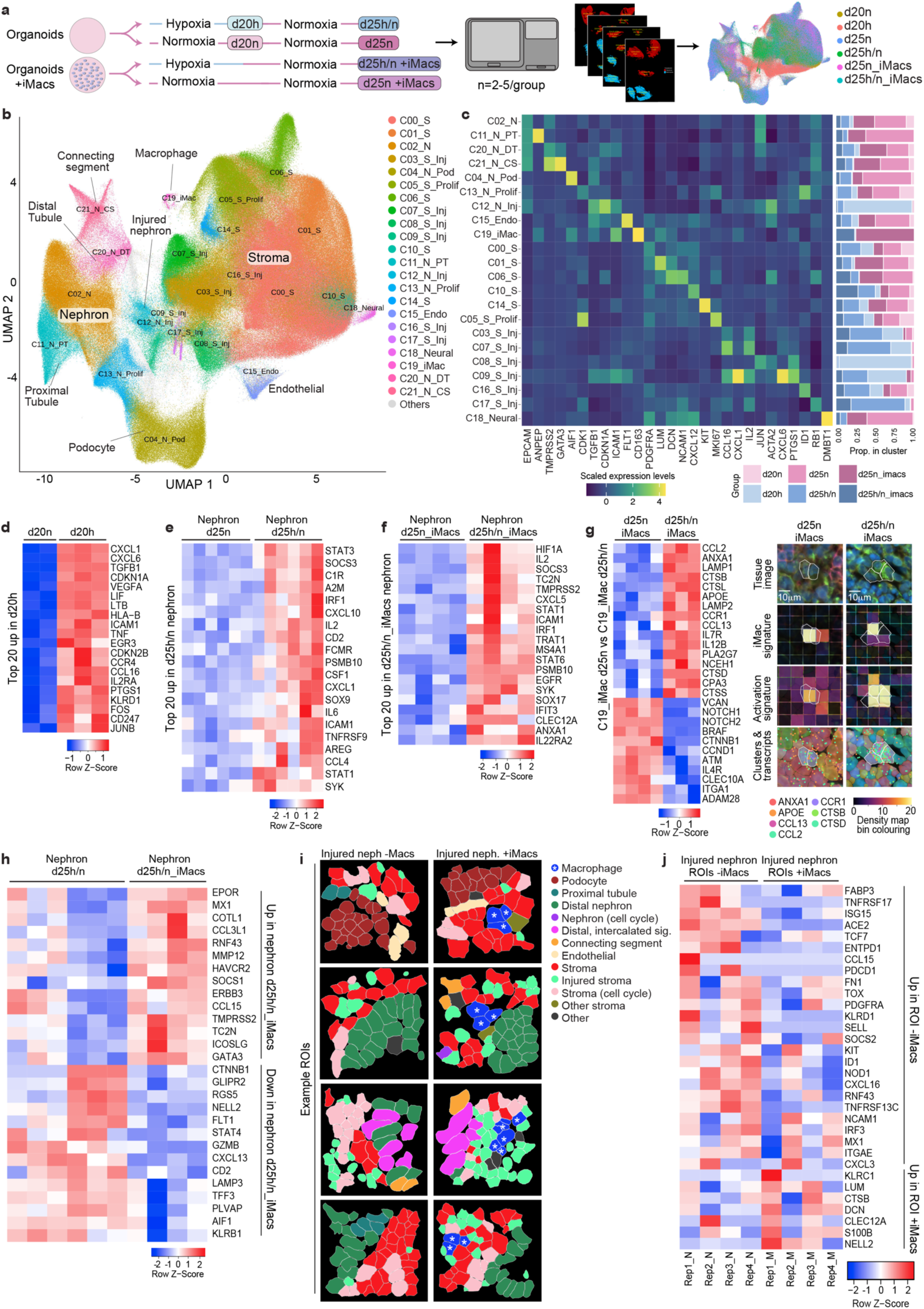
Spatial transcriptomic analysis of macrophage influence on injury and repair. (**a**) Experimental overview. (**b**) UMAP of clusters from Xenium spatial data. (**c**) Heatmap showing key marker genes supporting cluster identity (left) and proportions of cells within a cluster, coloured by sample group (right). (**d**) Heatmap of top 20 genes upregulated in d20h organoid spatial data compared to d20n supporting a hypoxic injury signature. (**e**) Heatmap of top 20 genes upregulated in the nephron lineage of d25h/n organoid spatial data compared to d25n supporting persistent injury and inflammation. (**f**) Heatmap of top 20 genes upregulated in the nephron lineage of d25h/n_iMacs organoid spatial data compared to d25n_iMacs, supporting persistent injury and inflammation. (**g**) Left - Heatmap of top up and downregulated genes in the macrophage cluster (C19) after hypoxic injury at d25, showing activation of inflammatory and phagocytic markers after injury. Right-representative regions of interest containing macrophages in d25n_iMacs and d25h_iMacs organoids showing tissue image (iMacs outlined), ‘iMac signature’ (density map for macrophage markers *CD14, CD68, PTPRC*), ‘activation signature’ (density map of select genes upregulated in injured organoids *ANXA1, APOE, CCL13, CCL2, CCR1, CTSB, CTSD*), cluster allocations and ‘activation signature’ transcripts. (**h**) Heatmap of the top differentially expressed genes between injured nephron clusters in organoids with or without iMacs. (**i**) Representative regions of interest (ROIs) containing injured nephrons, that have macrophages in the area, and comparable regions that lack macrophages. All ROIs are from d25h/n_imacs organoids. (**j**) Heatmap of differentially expressed genes upregulated in injured nephron regions containing or lacking macrophages. FDR and LogFC values available in Supplemental files 7 and 8. Columns show replicate samples in heatmaps d-h, columns show average of ROIs within a sample for j.

To validate expected hypoxic response signatures in this dataset, we performed differential expression analyses between sample groups at d20, which affirmed upregulation of genes related to hypoxic injury (Figure 8D). Similarly, comparison of nephron clusters in d25 injured and control organoids recovered expected signatures of tubular inflammation and ineffective repair in organoids with and without iMacs (Figure 8E-F, Supplemental File 7). Next, we focussed on how iMacs respond to hypoxia and their overall effect on nephron injury and repair in this model. Differential expression analysis of iMac clusters in d25 control and injured organoids supports a transition from a proliferative state engaged in tissue remodeling in normoxic conditions, to an activated inflammatory state with high lysosomal activity after hypoxia (Figure 8G, Table 3, Supplemental File 7).

**Table 3:**
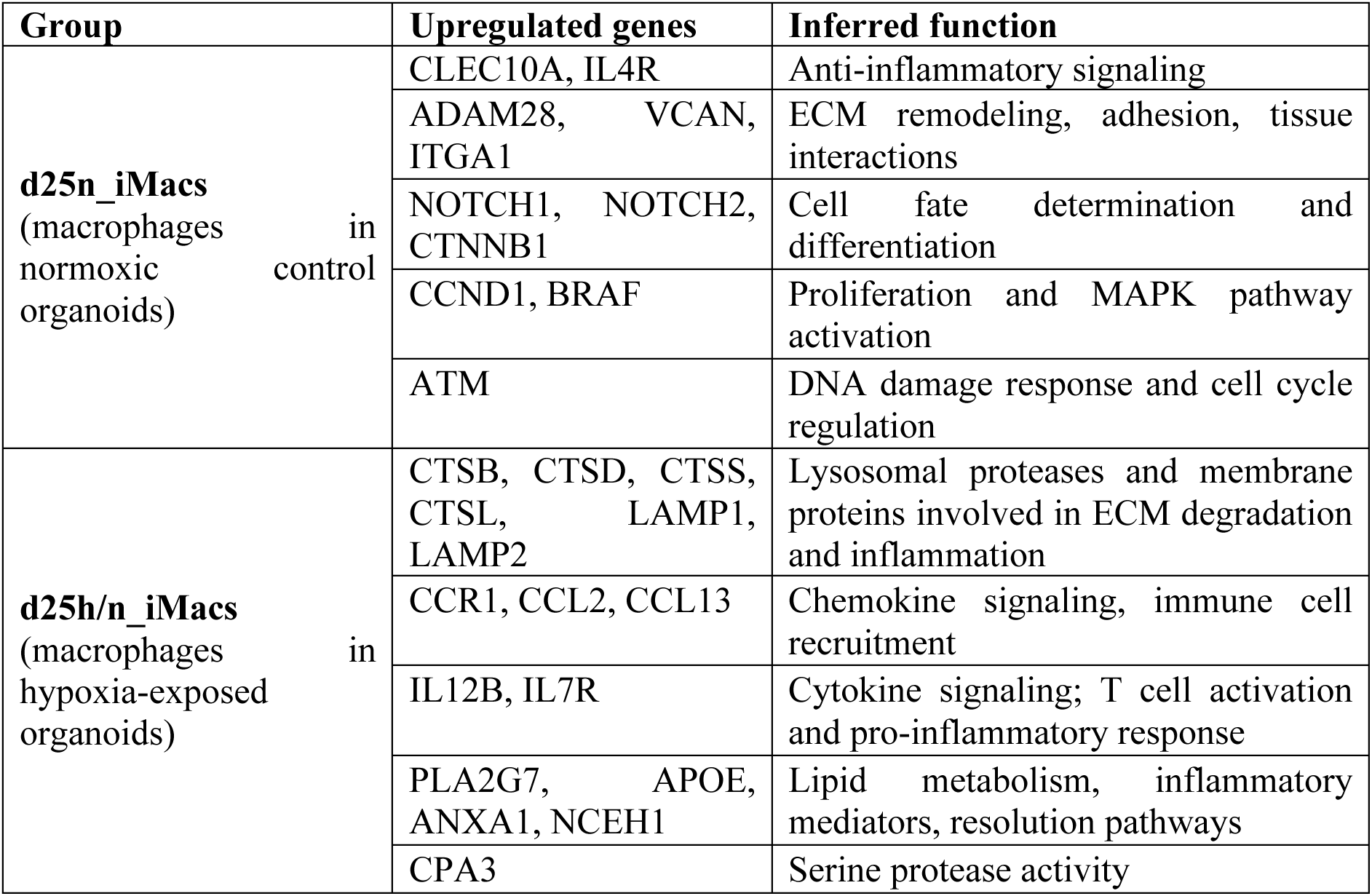
Gene expression changes and inferred functions in organoid-integrated macrophages after hypoxic injury.

Differential expression between nephron clusters in injured organoids +/- iMacs showed distinct injury responses. Nephrons in iMac-containing organoids expressed a broader range of ECM-remodelling (*SDC1, MMP12*), immune recruitment (*CCL15, CCL3L1, ICOSLG*), and immunomodulatory factors (*SOCS1, ICOSLG, MX1),* consistent with macrophage-mediated tissue remodelling after injury (Figure 8H). In contrast, injured nephrons from organoids lacking iMacs upregulated alternative immune recruitment signals (*CXCL13, CD2, KLRB1*), de-differentiation (*CTNNB1*), trafficking (*LAMP3*), and endothelial (*FLT1, PLVAP*) genes, suggesting an injury response involving intrinsic repair and vascular remodelling (Figure 8H).

To assess macrophage effects on the local environment, we compared injured nephron regions +/-iMacs, within d25h/n_iMac organoids (Figure 8I,J; Supplemental File 8). Regions containing injured nephrons and iMacs showed upregulation of macrophage activation, phagocytic and immunomodulatory genes, indicating an activated macrophage niche engaged in phagocytosis, cytokine signaling, and immune regulation. In contrast, injured regions lacking iMacs expressed genes associated with proliferation, survival, and repair programs (Supplemental File 8). Changes in the nephron epithelium within these regions identified a mixed profile spanning proteolysis and injury responses (*CTSB*), matrix organization and fibrotic regulation (*DCN, LUM*), immune inhibition and resolution (*KLRC1, CLEC12A*), and stress-related signalling (*S100B, NELL2*) in regions with iMacs (Figure 8J). Injured nephron regions lacking macrophages had similar, yet distinct signatures of fibrotic progression (*FN1, PDGFRA*), differentiation and stress response genes (*ID1, RNF43, FABP3, TOX*), and upregulation of injury markers (*CXCL3, CXCL16, CCL15*) (Figure 8J).

Limitations of this spatial analysis include the restricted scope of the targeted gene panel, variability in the plane of section across organoids, and extent of nephron loss after hypoxic injury. Nonetheless, in this kidney organoid model, iMacs respond to hypoxic injury by adopting an inflammatory, lysosome-active state. Gene expression signatures within these macrophages suggest they drive phagocytosis, ECM remodelling, fibrotic regulation and promote inflammation in the nephron epithelium.

## DISCUSSION

Ischemic AKI affects millions globally and frequently progresses to CKD, yet targeted therapies remain elusive due in part to limited human experimental models. Here we demonstrate that human kidney organoids recapitulate key molecular, metabolic and inflammatory signatures of ischemic AKI and maladaptive repair, establishing a scalable platform for mechanistic studies and therapeutic testing.

Our multi-omic analysis revealed striking conservation of disease signatures between organoids, animal models, and clinical AKI. Recent single cell and spatial profiling of over 18 human AKI biopsies identified injury and maladaptive repair signatures including elevated *CDKN1A* and *CCL2* expression and TGFB1, EGF, MAPK (FOS/JUN), NF-κB and JAK/STAT pathway signatures (18). We observed these same molecular programs in hypoxia-injured organoids, validating that human iPSC-derived kidney models preserve critical disease mechanisms. Validation of other established markers of injury and maladaptive repair including GDF15, S100A8/9, ICAM1, MMP7 and more, positions organoids as a human experimental model test new biomarkers, mechanisms and hypotheses emerging from clinical studies.

A particularly novel finding is the comprehensive metabolic dysregulation in injured organoids, mirroring CKD patient profiles. We observed dyslipidemia-like phenotypes with elevated oxysterols, sphingomyelins, and ceramides - lipid species associated with kidney pathology and oxidative stress in CKD patients (68, 74). Amino acid perturbations included reduced taurine, hypotaurine, tryptophan, and creatine, recapitulating similar deficiencies documented in CKD cohorts (69–72). Importantly, these metabolic signatures persisted despite five days of normoxic recovery, suggesting intrinsic metabolic memory that may drive progression. While previous organoid studies have focused primarily on transcriptional injury responses and select protein markers (21, 22, 75), our metabolomic profiling reveals a complementary layer of pathology and potential therapeutic targets.

Integration of iPSC-derived macrophages represents a significant advance in organoid complexity. Macrophages transitioned from a tissue-resident, remodeling phenotype in normoxia to an inflammatory, lysosome-active state after hypoxic injury, consistent with macrophage behaviour in murine AKI models and human biopsies (16, 17, 73). Spatial transcriptomics revealed that macrophage-containing injury niches expressed distinct gene programs compared to macrophage-depleted regions, highlighting localized immune-epithelial crosstalk. This builds substantially on prior organoid work by modeling immune cell contributions to injury outcomes - a critical gap given the central role of macrophages in determining repair trajectories.

Our results also illuminate injury-specific versus universal AKI mechanisms. Comparing our ischemic model with published cisplatin (21, 22) and hemolysis-induced (75) organoid studies reveals shared features (oxidative stress, injury marker induction, early fibrotic changes) alongside distinct signatures. Notably, we observed modest *HAVCR1/KIM1* induction despite scRNAseq supporting robust upregulation of injury markers within the proximal tubule (*GDF15, CCN1, ICAM1, IL32*). Additionally, where ineffective homology-directed DNA repair was identified as a key driver of cisplatin-induced maladaptive repair (22), this did not appear central to our ischemic model as *RAD51* and *FANCD2* expression recovered post-injury. Instead, hypoxia triggers initial injury and metabolic disruptions that evolve into sustained metabolic and inflammatory dysregulation. These findings underscore both shared and injury-specific mechanisms of AKI and highlight the importance of modelling distinct injury drivers to capture the mechanistic heterogeneity of clinical AKI.

Despite the expression of profibrotic factors including TGFB1, CCN2 and MMP7, and expansion of a stromal population expressing relevant markers, a strong fibrotic signature was not observed in this study. Limitations in the maturity of cell types within iPSC-derived kidney organoids along with their short lifespan and lack of functional vasculature, may affect the extent to which the fibrogenic process can be replicated *in vitro* (46). Future studies with alternative kidney organoid protocols enabling extended culture (22) or increased maturation of specific cell types (76–78) may enhance fibrotic progression in this model.

Because iPSC-derived kidney organoids resemble the human fetal kidney (29), this model may also inform links between intrauterine hypoxia and developmental programming of kidney disease (79). Our data show that hypoxia depletes nephron progenitor cells and causes loss of existing nephrons, offering a mechanistic explanation for reduced nephron endowment following fetal hypoxia.

In summary, we have established a human organoid model that recapitulates key mechanisms of ischemic AKI and early maladaptive repair through conserved inflammatory and metabolic programs. The integration of macrophages and spatial profiling reveals immune-epithelial interactions previously inaccessible in kidney organoid systems. This platform bridges the gap between animal models and clinical investigation, offering opportunities to validate patient-derived biomarkers, test therapeutic interventions, and dissect human-specific disease mechanisms. Future studies leveraging this model for drug screening, genetic perturbation, and patient-specific susceptibility will likely advance our understanding of AKI-to-CKD progression and accelerate therapeutic development.

## Disclosure

All authors declare no competing interests.

## Availability of data and materials

The datasets supporting the conclusions of this study will be made available on publication. Bulk, single cell, and spatial transcriptomic data have been deposited in the National Center for Biotechnology Information’s GEO database (bulk accession No. GSE236379; bulk iMacs accession No. GSE278562; scRNAseq accession No. GSE242426; spatial accession No. GSE307588). scRNAseq quality control and analysis, an interactive marker visualisation website, and corresponding code are stored and documented in a GitHub repository https://github.com/MonashBioinformaticsPlatform/sc-hyp-org. Proteomic data are available via ProteomeXchange with identifier PXD042882, metabolomic data are available at https://store.erc.monash.edu.au/experiment/view/16985/. The three iPSC lines in this study have been used under material transfer agreements with the Commonwealth Scientific and Industrial Research Organisation (CSIRO) and Murdoch Children’s Research Institute (MCRI).

## Acknowledgements

The authors acknowledge use of the MHTP Medical Genomics Facility, Micromon Genomics, Monash Histology Platform, Monash Bioinformatics Platform, and Monash Micro Imaging. This study used infrastructure enabled by Bioplatforms Australia and the National Collaborative Research Infrastructure Strategy at the Monash Proteomics and Metabolomics Platform. We gratefully acknowledge the WEHI Spatial Omics platform laboratory for conducting the Xenium experiments, Anne Hempel for performing the HIF1A Western Blot and Rachel Lam for early contributions to the project. We thank Sara Howden and Melissa Little for support and providing the 522 and 808 iPSC lines, and Andrew Laslett for the D4C4 iPSC line.

## Author contributions

A.B.N.N, D.J.N.P. and A.N.C. conceived and designed the study; A.B.N.N performed all organoid experiments, curated and interpreted data; M.P. performed and generated figures for deconvolution and bulk RNAseq analyses; J.R.S. and H.L. prepared samples for proteomics, generated and analysed data; J.L.M performed and analysed ICAM and KIM1 staining; C.K.B. undertook the untargeted metabolomics including sample preparation, LC-MS analysis and data analysis; R.B.S. provided resources, methodology and supervision for the MS study. C.A.W. and Z.C. provided iPSC-derived human macrophages.

Y.L. and W.S. performed the spatial transcriptomic analysis, with interpretation and input from A.N.C., under the supervision of W.S.; Funding was acquired by A.N.C; The manuscript was authored and revised by A.B.N.N. and A.N.C., with contributions from all authors; The project was supervised by A.N.C and D.J.N.P. All authors subsequently reviewed and approved the manuscript.

## SUPPLEMENTAL MATERIALS

Supplemental Figures 1 to 8

Supplemental Tables 1 to 2

Supplemental Files 1 to 8

## References

1. Abebe A, Kumela K, Belay M, Kebede B, Wobie Y. Mortality and predictors of acute kidney injury in adults: a hospital-based prospective observational study. Scientific Reports. 2021;11(1):15672.

2. Mehta RL, Cerdá J, Burdmann EA, Tonelli M, García-García G, Jha V, et al. International Society of Nephrology’s 0by25 initiative for acute kidney injury (zero preventable deaths by 2025): a human rights case for nephrology. The Lancet. 2015;385(9987):2616–43.

3. Molitoris BA. Low-Flow Acute Kidney Injury: The Pathophysiology of Prerenal Azotemia, Abdominal Compartment Syndrome, and Obstructive Uropathy. Clinical Journal of the American Society of Nephrology. 2022;17(7).

4. O’Neal JB, Shaw AD, Billings FTt. Acute kidney injury following cardiac surgery: current understanding and future directions. Crit Care. 2016;20(1):187.

5. Bonventre JV, Yang L. Cellular pathophysiology of ischemic acute kidney injury. J Clin Invest. 2011;121(11):4210–21.

6. Bendall AC, See EJ, Toussaint ND, Fazio T, Tan SJ. Community-acquired versus hospital-acquired acute kidney injury at a large Australian metropolitan quaternary referral centre: incidence, associations and outcomes. Intern Med J. 2023;53(8):1366–75.

7. Bhargava P, Schnellmann RG. Mitochondrial energetics in the kidney. Nat Rev Nephrol. 2017;13(10):629–46.

8. Kusaba T, Lalli M, Kramann R, Kobayashi A, Humphreys BD. Differentiated kidney epithelial cells repair injured proximal tubule. Proceedings of the National Academy of Sciences. 2014;111(4):1527–32.

9. Kumar S, Liu J, Pang P, Krautzberger AM, Reginensi A, Akiyama H, et al. Sox9 Activation Highlights a Cellular Pathway of Renal Repair in the Acutely Injured Mammalian Kidney. Cell Reports. 2015;12(8):1325–38.

10. Kang HM, Huang S, Reidy K, Han SH, Chinga F, Susztak K. Sox9-Positive Progenitor Cells Play a Key Role in Renal Tubule Epithelial Regeneration in Mice. Cell Rep. 2016;14(4):861–71.

11. Jin Y, Ratnam K, Chuang PY, Fan Y, Zhong Y, Dai Y, et al. A systems approach identifies HIPK2 as a key regulator of kidney fibrosis. Nat Med. 2012;18(4):580–8.

12. Gerhardt LMS, Koppitch K, van Gestel J, Guo J, Cho S, Wu H, et al. Lineage Tracing and Single-Nucleus Multiomics Reveal Novel Features of Adaptive and Maladaptive Repair after Acute Kidney Injury. Journal of the American Society of Nephrology. 2023;34(4).

13. Li Z, Liu Z, Luo M, Li X, Chen H, Gong S, et al. The pathological role of damaged organelles in renal tubular epithelial cells in the progression of acute kidney injury. Cell Death Discovery. 2022;8(1):239.

14. Kang HM, Ahn SH, Choi P, Ko Y-A, Han SH, Chinga F, et al. Defective fatty acid oxidation in renal tubular epithelial cells has a key role in kidney fibrosis development. Nature Medicine. 2015;21(1):37–46.

15. Sugahara M, Tanaka S, Tanaka T, Saito H, Ishimoto Y, Wakashima T, et al. Prolyl Hydroxylase Domain Inhibitor Protects against Metabolic Disorders and Associated Kidney Disease in Obese Type 2 Diabetic Mice. Journal of the American Society of Nephrology. 2020;31(3).

16. Munro DAD, Hughes J. The Origins and Functions of Tissue-Resident Macrophages in Kidney Development. Front Physiol. 2017;8:837.

17. Li Z, Zimmerman KA, Yoder BK. Resident Macrophages in Cystic Kidney Disease. Kidney360. 2021;2(1):167-75.

18. Lake BB, Menon R, Winfree S, Hu Q, Ferreira RM, Kalhor K, et al. An atlas of healthy and injured cell states and niches in the human kidney. Nature. 2023;619(7970):585–94.

19. Freedman BS, Brooks CR, Lam AQ, Fu H, Morizane R, Agrawal V, et al. Modelling kidney disease with CRISPR-mutant kidney organoids derived from human pluripotent epiblast spheroids. Nature Communications. 2015;6(1):8715.

20. Nunez-Nescolarde AB, Nikolic-Paterson DJ, Combes AN. Human Kidney Organoids and Tubuloids as Models of Complex Kidney Disease. The American Journal of Pathology. 2022;192(5):738–49.

21. Digby JLM, Vanichapol T, Przepiorski A, Davidson AJ, Sander V. Evaluation of cisplatin-induced injury in human kidney organoids. Am J Physiol Renal Physiol. 2020;318(4):F971–F8.

22. Gupta N, Matsumoto T, Hiratsuka K, Garcia Saiz E, Galichon P, Miyoshi T, et al. Modeling injury and repair in kidney organoids reveals that homologous recombination governs tubular intrinsic repair. Sci Transl Med. 2022;14(634):eabj4772.

23. Ohmori T, De S, Tanigawa S, Miike K, Islam M, Soga M, et al. Impaired NEPHRIN localization in kidney organoids derived from nephrotic patient iPS cells. Scientific Reports. 2021;11(1):3982.

24. Takasato M, Er PX, Chiu HS, Maier B, Baillie GJ, Ferguson C, et al. Kidney organoids from human iPS cells contain multiple lineages and model human nephrogenesis. Nature. 2015;526(7574):564–8.

25. Taguchi A, Kaku Y, Ohmori T, Sharmin S, Ogawa M, Sasaki H, et al. Redefining the in vivo origin of metanephric nephron progenitors enables generation of complex kidney structures from pluripotent stem cells. Cell Stem Cell. 2014;14(1):53–67.

26. Morizane R, Lam AQ, Freedman BS, Kishi S, Valerius MT, Bonventre JV. Nephron organoids derived from human pluripotent stem cells model kidney development and injury. Nature Biotechnology. 2015;33(11):1193–200.

27. Howden SE, Little MH. Generating Kidney Organoids from Human Pluripotent Stem Cells Using Defined Conditions. Methods Mol Biol. 2020;2155:183–92.

28. Takasato M, Er PX, Becroft M, Vanslambrouck JM, Stanley EG, Elefanty AG, et al. Directing human embryonic stem cell differentiation towards a renal lineage generates a self-organizing kidney. Nature Cell Biology. 2014;16(1):118–26.

29. Takasato M, Er PX, Chiu HS, Maier B, Baillie GJ, Ferguson C, et al. Kidney organoids from human iPS cells contain multiple lineages and model human nephrogenesis. Nature. 2015;526(7574):564–8.

30. Berrocal-Rubio MA, Pawer YDJ, Dinevska M, De Paoli-Iseppi R, Widodo SS, Gleeson J, et al. Discovery of NRG1-VII: the myeloid-derived class of NRG1. BMC Genomics. 2024;25(1):814.

31. Rajab N, Angel PW, Deng Y, Gu J, Jameson V, Kurowska-Stolarska M, et al. An integrated analysis of human myeloid cells identifies gaps in in vitro models of in vivo biology. Stem Cell Reports. 2021;16(6):1629–43.

32. Grubman A, Choo XY, Chew G, Ouyang JF, Sun G, Croft NP, et al. Transcriptional signature in microglia associated with Aβ plaque phagocytosis. Nature Communications. 2021;12(1):3015.

33. Ritchie ME, Phipson B, Wu D, Hu Y, Law CW, Shi W, et al. limma powers differential expression analyses for RNA-sequencing and microarray studies. Nucleic acids research. 2015;43(7):e47-e.

34. Law CW, Chen Y, Shi W, Smyth GK. voom: Precision weights unlock linear model analysis tools for RNA-seq read counts. Genome biology. 2014;15(2):1–17.

35. Powell D. Degust: interactive RNA-seq analysis. Drpowell/Degust. 2015;4(1):4.1.

36. Sherman BT, Hao M, Qiu J, Jiao X, Baseler MW, Lane HC, et al. DAVID: a web server for functional enrichment analysis and functional annotation of gene lists (2021 update). Nucleic acids research. 2022;50(W1):W216–W21.

37. Subramanian A, Tamayo P, Mootha VK, Mukherjee S, Ebert BL, Gillette MA, et al. Gene set enrichment analysis: a knowledge-based approach for interpreting genome-wide expression profiles. Proceedings of the National Academy of Sciences. 2005;102(43):15545–50.

38. Howden SE, Wilson SB, Groenewegen E, Starks L, Forbes TA, Tan KS, et al. Plasticity of distal nephron epithelia from human kidney organoids enables the induction of ureteric tip and stalk. Cell Stem Cell. 2021;28(4):671–84.e6.

39. Hao Y, Hao S, Andersen-Nissen E, Mauck WM, 3rd, Zheng S, Butler A, et al. Integrated analysis of multimodal single-cell data. Cell. 2021;184(13):3573–87 e29.

40. Teo GC, Polasky DA, Yu F, Nesvizhskii AI. Fast Deisotoping Algorithm and Its Implementation in the MSFragger Search Engine. J Proteome Res. 2021;20(1):498–505.

41. Shah AD, Goode RJA, Huang C, Powell DR, Schittenhelm RB. LFQ-Analyst: An Easy-To-Use Interactive Web Platform To Analyze and Visualize Label-Free Proteomics Data Preprocessed with MaxQuant. Journal of Proteome Research. 2020;19(1):204–11.

42. Cable DM, Murray E, Zou LS, Goeva A, Macosko EZ, Chen F, et al. Robust decomposition of cell type mixtures in spatial transcriptomics. Nat Biotechnol. 2022;40(4):517–26.

43. Law CW, Chen Y, Shi W, Smyth GK. voom: Precision weights unlock linear model analysis tools for RNA-seq read counts. Genome Biol. 2014;15(2):R29.

44. Ritchie ME, Phipson B, Wu D, Hu Y, Law CW, Shi W, et al. limma powers differential expression analyses for RNA-sequencing and microarray studies. Nucleic Acids Res. 2015;43(7):e47.

45. Smyth GK. Linear models and empirical bayes methods for assessing differential expression in microarray experiments. Stat Appl Genet Mol Biol. 2004;3:Article3.

46. Combes AN, Phipson B, Lawlor KT, Dorison A, Patrick R, Zappia L, et al. Single cell analysis of the developing mouse kidney provides deeper insight into marker gene expression and ligand-receptor crosstalk. Development. 2019;146(12):dev178673.

47. Majmundar AJ, Wong WJ, Simon MC. Hypoxia-Inducible Factors and the Response to Hypoxic Stress. Molecular Cell. 2010;40(2):294–309.

48. Venkatachalam MA, Weinberg JM, Kriz W, Bidani AK. Failed Tubule Recovery, AKI-CKD Transition, and Kidney Disease Progression. Journal of the American Society of Nephrology. 2015;26(8):1765.

49. Koshiji M, Kageyama Y, Pete EA, Horikawa I, Barrett JC, Huang LE. HIF-1alpha induces cell cycle arrest by functionally counteracting Myc. Embo j. 2004;23(9):1949–56.

50. Fontecha-Barriuso M, Martin-Sanchez D, Martinez-Moreno JM, Monsalve M, Ramos AM, Sanchez-Niño MD, et al. The Role of PGC-1α and Mitochondrial Biogenesis in Kidney Diseases. Biomolecules [Internet]. 2020; 10(2).

51. Basile DP, Anderson MD, Sutton TA. Pathophysiology of acute kidney injury. Compr Physiol. 2012;2(2):1303–53.

52. Zager RA, Johnson ACM. Acute kidney injury induces dramatic p21 upregulation via a novel, glucocorticoid-activated, pathway. Am J Physiol Renal Physiol. 2019;316(4):F674–f81.

53. Bülow RD, Boor P. Extracellular Matrix in Kidney Fibrosis: More Than Just a Scaffold. Journal of Histochemistry & Cytochemistry. 2019;67(9):643–61.

54. Humphreys BD. Mechanisms of Renal Fibrosis. Annu Rev Physiol. 2018;80:309–26.

55. Ferenbach DA, Bonventre JV. Mechanisms of maladaptive repair after AKI leading to accelerated kidney ageing and CKD. Nat Rev Nephrol. 2015;11(5):264–76.

56. Liu J, Kumar S, Heinzel A, Gao M, Guo J, Alvarado GF, et al. Renoprotective and Immunomodulatory Effects of GDF15 following AKI Invoked by Ischemia-Reperfusion Injury. J Am Soc Nephrol. 2020;31(4):701–15.

57. Nair V, Robinson-Cohen C, Smith MR, Bellovich KA, Bhat ZY, Bobadilla M, et al. Growth Differentiation Factor-15 and Risk of CKD Progression. J Am Soc Nephrol. 2017;28(7):2233–40.

58. Cangul H. Hypoxia upregulates the expression of the NDRG1 gene leading to its overexpression in various human cancers. BMC Genet. 2004;5:27.

59. Higgins DF, Biju MP, Akai Y, Wutz A, Johnson RS, Haase VH. Hypoxic induction of Ctgf is directly mediated by Hif-1. American Journal of Physiology-Renal Physiology. 2004;287(6):F1223–F32.

60. Combes AN, Zappia L, Er PX, Oshlack A, Little MH. Single-cell analysis reveals congruence between kidney organoids and human fetal kidney. Genome Med. 2019;11(1):3.

61. Wilson SB, Howden SE, Vanslambrouck JM, Dorison A, Alquicira-Hernandez J, Powell JE, et al. DevKidCC allows for robust classification and direct comparisons of kidney organoid datasets. Genome Med. 2022;14(1):19.

62. Phipson B, Sim CB, Porrello ER, Hewitt AW, Powell J, Oshlack A. propeller: testing for differences in cell type proportions in single cell data. Bioinformatics. 2022;38(20):4720–6.

63. Wang W-g, Sun W-x, Gao B-s, Lian X, Zhou H-l. Cell Cycle Arrest as a Therapeutic Target of Acute Kidney Injury. Current Protein & Peptide Science. 2017;18(12):1224–31.

64. Taguchi K, Elias BC, Sugahara S, Sant S, Freedman BS, Waikar SS, et al. Cyclin G1 induces maladaptive proximal tubule cell dedifferentiation and renal fibrosis through CDK5 activation. J Clin Invest. 2022;132(23).

65. Wheaton WW, Chandel NS. Hypoxia. 2. Hypoxia regulates cellular metabolism. Am J Physiol Cell Physiol. 2011;300(3):C385–93.

66. Rodriguez D, Watts D, Gaete D, Sormendi S, Wielockx B. Hypoxia Pathway Proteins and Their Impact on the Blood Vasculature. International Journal of Molecular Sciences [Internet]. 2021; 22(17).

67. Siems W, Quast S, Peter D, Augustin W, Carluccio F, Grune T, et al. Oxysterols are increased in plasma of end-stage renal disease patients. Kidney Blood Press Res. 2005;28(5-6):302-6.

68. Nicholson RJ, Pezzolesi MG, Summers SA. Rotten to the Cortex: Ceramide-Mediated Lipotoxicity in Diabetic Kidney Disease. Front Endocrinol (Lausanne). 2020;11:622692.

69. Chen D-Q, Cao G, Chen H, Argyopoulos CP, Yu H, Su W, et al. Identification of serum metabolites associating with chronic kidney disease progression and anti-fibrotic effect of 5-methoxytryptophan. Nature Communications. 2019;10(1):1476.

70. Cheng Y, Li Y, Benkowitz P, Lamina C, Köttgen A, Sekula P. The relationship between blood metabolites of the tryptophan pathway and kidney function: a bidirectional Mendelian randomization analysis. Scientific Reports. 2020;10(1):12675.

71. Salazar JH. Overview of Urea and Creatinine. Laboratory Medicine. 2014;45(1):e19–e20.

72. Post A, Tsikas D, Bakker SJL. Creatine is a Conditionally Essential Nutrient in Chronic Kidney Disease: A Hypothesis and Narrative Literature Review. Nutrients. 2019;11(5).

73. Polonsky M, Gerhardt LMS, Yun J, Koppitch K, Colon KL, Amrhein H, et al. Spatial transcriptomics defines injury specific microenvironments and cellular interactions in kidney regeneration and disease. Nat Commun. 2024;15(1):7010.

74. Florens N, Calzada C, Lyasko E, Juillard L, Soulage CO. Modified Lipids and Lipoproteins in Chronic Kidney Disease: A New Class of Uremic Toxins. Toxins (Basel). 2016;8(12).

75. Przepiorski A, Vanichapol T, Espiritu EB, Crunk AE, Parasky E, McDaniels MD, et al. Modeling oxidative injury response in human kidney organoids. Stem Cell Res Ther. 2022;13(1):76.

76. Nunez-Nescolarde AB, Nikolic-Paterson DJ, Combes AN. Human Kidney Organoids and Tubuloids as Models of Complex Kidney Disease. The American Journal of Pathology. 2022.

77. Vanslambrouck JM, Wilson SB, Tan KS, Groenewegen E, Rudraraju R, Neil J, et al. Enhanced metanephric specification to functional proximal tubule enables toxicity screening and infectious disease modelling in kidney organoids. Nature Communications. 2022;13(1):5943.

78. Little MH, Combes AN. Kidney organoids: accurate models or fortunate accidents. Genes Dev. 2019;33(19-20):1319–45.

79. Luyckx VA, Brenner BM. Clinical consequences of developmental programming of low nephron number. The Anatomical Record. 2020;303(10):2613–31.

