## Supplemental Materials for "Hypoxic injury triggers maladaptive repair in human kidney organoids"

Nunez-Nescolarde et al.

**This file includes**:

Supplemental Materials and Methods

Supplemental Figures 1 to 8

Supplemental Tables 1 to 2

#### Supplemental Materials and Methods

##### Generation of Kidney Organoids

Human iPSCs were directed to differentiate into kidney organoids using the Takasato protocol (1-3), with minor modifications. iPSCs were seeded on Matrigel-coated 6-well plates (Corning, cat no. FAL35427) using E8 media (Thermo Fisher Scientific) supplemented with 10 µM of ROCK inhibitor Y-27632 (In Vitro Technologies) and 1% Pen/Strep (Thermo Fisher Scientific). The next day (day 0 of differentiation, d0), media was changed to Essential 6 (E6) (Thermo Fisher Scientific), which was initially supplemented with CHIR99021 (In Vitro Technologies) and 1% Pen/Strep (Thermo Fisher Scientific). On day 4, 200 ng/ml of FGF9 (In Vitro Technologies) and 1 µg/ml of Heparin (Sigma-Aldrich) were added to E6. On d7, cells were dissociated with TrypLE Select Enzyme (1X) (Thermo Fisher Scientific) and 150 000 cells were centrifuged in 1.5 ml tubes to generate each organoid. Newly formed organoids were treated with CHIR99021 in E6 medium for one hour (Supplemental Table S1). Organoids were cultured in E6 supplemented with 200 ng/ml of FGF9, 1 µg/ml of Heparin and 1 % Pen/Strep until d13. After, only E6 and Pen/Strep were used.

Cell seeding density and CHIR99021 concentrations were optimized for each iPSC line as follows. 522.3 and 808.5 lines were seeded at 50,000-80,000 cells per well at d-1, treated with 4 µM CHIR99021 from d0-d4. The D4C4 line was seeded at 70,000-100,000 at d-1 and treated with 6 µM CHIR99021 from d0-d4. Differentiations from all lines were dissociated into single cell suspensions at d7 and 150,000 cells were aggregated by centrifugation to generate organoids. Concentrations of CHIR99021 used to induce nephron formation at d7 were 4 µM (522.3 and 808.5) and 5 µM (D4C4).

##### Kidney Organoid-Macrophage Co-culture

iPSC-derived macrophages were differentiated from dual reporter Cln29.2 OAS2:mCherry/Cln109.11 REL:YFP PB001.1 human iPSCs as described (4, 5). Briefly, iPSCs were maintained in Essential 8 medium (Thermo Fisher, #A1517001) on Matrigel-coated dishes and passaged at 70–80% confluence. On Day 0, cells were dissociated with 0.5 mM EDTA, collected by centrifugation (200 g, 4 min), and 1–1.5 × $10^6$ cells were replated in “MAGIC” medium for embryoid body (EB) formation in non-coated 10 cm suspension dishes (Corning). EBs formed overnight and were cultured for up to 4 weeks with media changes on Days 1, 2, 4, 8, 11, 15, and Week 3, using growth factor supplements reported in (4). During medium changes, EBs were transferred gently using a 10 ml serological pipette, allowed to settle by gravity (~10 min), and resuspended slowly in fresh medium. On Day 11, EBs were allowed to settle (<6 min) and the progenitor-containing supernatant was collected, centrifuged (350 g, 4 min), and the pellet was resuspended and counted. Cells were plated at 1 × $10^5$ cells/well in a 6-well plate in RPMI + 10% FBS + 100 ng/ml M-CSF-1

(PrepoTech cat. No. 300-25) for macrophage maturation. Macrophages matured after 4 days, with medium changed every 3 days for extended culture. Macrophages were then dissociated with 0.5mM EDTA and enriched by human CD14 microbeads (Miltenyi Biotec) with a purity of CD14 expression higher than 95% tested by flow cytometry. Integration of macrophages was performed on day 7 of kidney organoid differentiation. After single cell dissociation with TrypLE Select Enzyme (1X) (Thermo Fisher Scientific), 7500 macrophages were incorporated into each microtube, which already contained 142500 cells from the kidney differentiation, representing 5% macrophages: 95% kidney progenitors at day 7. After mixing with a pipette, each 3D organoid was formed by centrifugation and transferred to a transwell membrane for culture as previously described.

**Hypoxic Injury**

iPSCs and organoids were cultured in standard conditions in a humidified incubator (37C, 5% $CO_2$, 19% $O_2$, 76% $N_2$) until day 18 of differentiation. Control organoids were maintained at 19% $O_2$ from this point, while experimental groups were cultured in a humidified incubator under hypoxic incubators (37C, 5% $CO_2$, 1% $O_2$, 94% $N_2$) for 48h then harvested immediately (d20) or after a 5 day recovery at 19% $O_2$ (d25). For bulk RNA-sequencing, quantitative PCR, proteomics and metabolomics studies, organoids were snap-frozen, for immunostaining and spatial transcriptomics, organoids were fixed in 4% paraformaldehyde (PFA), at d20 and d25.

**Quantitative PCR**

RNA isolation and purification was performed with the Isolate II RNA Mini Kit (Bioline, cat no. BIO-52072), following the manufacturer's instructions. RNA quantification was carried out using NanoDrop ND-1000 (ThermoFisher). cDNA was synthetized using the QuantiTect Rev. Transcription kit (QIAGEN, cat no. 205311). Quantitative PCR was performed using the Qiagen Quantinova SYBR green (Qiagen, cat no. 208054) on a 384-well Biorad unit. Gene expression was calculated using ΔCT method and beta-actin (*ACTB)* for normalization. Error bars represent standard deviation (SD) of biological replicates of the three cell lines. Data is expressed as mean ± SD and analyzed by analysis of variance (ANOVA) with Tukey's multiple comparisons test. Differences were considered significant at *P < 0.05, **P < 0.01, ***P < 0.001.

Primers used were: HIF1A (F: CATAAAGTCTGCAACATGGAAGGT, R: TTGATGGGTGAGGAATGGGTT), VEGFA (F: TGCAGATTATGCGGATCAAACC, R: TGCATTCACATTTGTTGTGCTGTAG), HAVCR1 (KIM-1) (F: CTGCAGGGAGCAATAAGGAG, R: TCCAAAGGCCATCTGAAGAC), HK1 (F: GAGTTTGACCTGGATGTGGTTGC, R: CCTCCATGTAGCAGGCATTGCT), TGFB1 (F:

TACCTGAACCCGTGTTGCTCTC, R: GTTGCTGAGGTATCGCCAGGAA), COL1A1 (F: TCTGCGACAACGGCAAGGTG, R: GACGCCGGTGGTTTCTTGGT), FN1 (F: ACAACACCGAGGTGACTGAGAC, R: GGACACAACGATGCTTCCTGAG), CCL2 (F: AGGTGACTGGGGCATTGAT, R: GCCTCCAGCATGAAAGTCTC), CDKN1A (F: AGGGGACAGCAGAGGAAG, R: GCGTTTGGAGTGGTAGAAATCTG), NANOG (F: AATACCTCAGCCTCCAGCAGATG, R: TGCGTCACACCATTGCTATTCTTC), TBX6 (F: CATCCACGAGAATTGTACCCG, R: AGCAATCCAGTTTAGGGGTGT), T (F: AGGTACCCAACCCTGAGGA, R: GCAGGTGAGTTGTCAGAATAGGT), SIX2 (F: CTGGAGAGCCACCAGTTCTC, R: GCTGCGACTCTTTTCCTTGA), OSR1 (F: CCTTCCTTCAGGCAGTGAAC, R: CGGCACTTTGGAGAAAGAAG), EYA1 (F: ATCTAACCAGCCCGCATAGC, R: GTGCCATTGGGAGTCATGGA), LHX1 (F: ATGCAACCTGACCGAGAAGT, R: CAGGTCGCTAGGGGAGATG), GATA3 (F: GCCCCTCATTAAGCCCAAG, R: TTGTGGTGGTCTGACAGTTCG), SLC12A1 (F: AGTGCCCAGTAATACCAATCGC, R: GCCTAAAGCTGATTCTGAGTCTT), UMOD (F: TGACCAACTGCTATGCCACACC, R: GAACATCTGGACGGAAAATCGGC), LRP2 (F: AAATTGAGCACAGCACCTTTGA, R: TCTGCTTTCCTGACTCGAATAATG), CALB1 (F: TCCAGGGAATCAAAATGTGTGG, R: GCACAGATCCTTCAGTAAAGCA), NPHS1 (F: AGTGTGGCTAAGGGATTACCC, R: TCACCGTGAATGTTCTGTTCC), PECAM1 (F: GCAGCATCGTGGTCAACATAA, R: GCAGGACAGGTTCAGTCTTTCA), GDF15 (F: GTTAGCCAAAGACTGCCACTG, R: CCTTGAGCCCATTCCACA), PPARGC1A (F: GTCACCACCCAAATCCTTAT, R: ATCTACTGCCTGGAGACCTT), ACTB (F: GTTACAGGAAGTCCCTTGCCATCC, R: CACCTCCCCTGTGTGGACTTGGG).

**Western blot**

The nuclear fraction of pools of 3 organoids subjected to 24 h and 48 h of hypoxia was used for Western blot analysis. Organoids were sonicated and lysed in 0.4 ml of RIPA lysis buffer. Samples were centrifuged at 875 g at 4°C for 30 min to remove cell debris. Protein concentrations were measured using a modified Lowry protein assay with BSA as the protein standard (DC Protein Assay Kit, Bio-Rad). Extracted proteins were separated by electrophoresis on a 10% SDS–PAGE gel and then transferred to a PVDF membrane. After blocking for 30 min at 4°C in blocking buffer (5% skim milk powder in PBS with 0.1% Tween-20), the membrane was incubated overnight with a mouse monoclonal anti-HIF-1α primary antibody (1:300, Novus Biologicals, cat. No. NB100-105). The membrane was then washed and incubated for 30 min at room temperature with a goat anti-mouse HRP secondary antibody (1:5000, Novus Biologicals, cat. No. HAF007). β-actin was used as a loading control (1:5000, Novus Biologicals, NB600). Immunoreactive bands were detected using the ChemiDoc MP Imager (Bio-Rad).

**Bulk RNA-seq and Data Analysis**

Total RNA was isolated from kidney organoids on d20 and d25 of differentiation in triplicates using the Isolate II RNA Mini Kit (Bioline, cat no. BIO-52072). Sequencing was performed at MHTP Medical Genomics at Monash University using the 3'-Multiplex RNA-seq method, an in-house developed method that captures and sequences the 3'end of pA transcripts, suitable for multiplexing. RNA integrity was assessed with Bioanalyzer (Agilet Technologies) (see Supplemental Table 7 for RIN scores). All samples were processed starting with 50 ng of total RNA. Libraries were made by individual first strand synthesis to add the i7 index sequences indicated in Supplemental Table 3. Library construction of pooled cDNA (tagged with i7 indexes) was completed by adding the full P7 and P5 arms by tagmentation and PCR. Quality control (QC) for the library pool included analysis concentration and quality using a Qubit, Bioanalyzer and qPCR. It had a mean size of 466bp at a concentration of 7.1 nM. The library pool was loaded at 1000pM (based upon size adjusted qPCR) for on-board denaturation and clustering which resulted in a loading metric of 99.7 and 90% >Q30. 463.2 Million reads were assigned to indexes. Numbers of reads assigned to each sample are shown in Supplemental Table 3**.** Libraries were run on Illumina NextSeq 2000.

Sequencing data were indexed to the human GRCh38 reference genome using the STAR alignment tool v.2.7.10 and human Ensembl GTF annotation file v.106 (6). Subsequently, fastq files were aligned to the indexed genome to generate BAM files. The BAM files were then sorted and indexed using "samtools" software v.1.15.1 (7). To eliminate duplicated reads based on Unique Molecular Identifiers (UMIs), the "umi_tools" software v.1.1.2 was employed (8). Count data was generated using "featureCounts" tool v.2.0.1 on sorted deduplicated BAM files (9). The resulting count data were imported into the R statistical programming environment to assemble the gene expression matrix. "biomaRt" R package v.2.46.3 was used to map Ensembl gene Ids to gene symbols (10). Data were normalized using the CPM method and differentially expressed genes were recognized using voom estimated weights and limma R package (11, 12) and Degust software (13) (BH adj-p-value < 0.05 & absolute LogFC > 0.5).

Differentially expressed genes were used to perform enrichment analysis using the DAVID online tool (14). Further analysis was performed with clusterProfiler R package v.4.6.1, taking ranked symbols based on LogFCs. For GSEA analysis, normalized gene expression profiles were imported into GSEA tool v.4.2.3 (15) employing the MSigDB database (16).

**Single cell RNA-seq and Data Analysis**

12 samples from iPSC-derived kidney organoids were labelled with Cell Plex (10x Genomics) Cell Modified Oligonucleotides (CMO) for multiplexing: n1_d20 (CMO301), n2_d20 (CMO302), n3_d20 (CMO303), h1_d20 (CMO304), h2_d20 (CMO305), h3_d20 (CMO306), n1_d25 (CMO307), n2_d25 (CMO308), n3_d25 (CMO309), h1_d25 (CMO310), h2_d25 (CMO311), and h3_d25 (CMO312). These 12 samples correspond to 4 conditions with 3 replicates per condition and a pool of 3 organoids for each replicate. Libraries were prepared using 10X Genomics Chromium Single Cell 5' V3.1 chemistry, for GEX and FB/CellPlex libraries (10X Genomics; San Francisco CA, USA), according to the manufacturer's instructions and sequenced on MGITech MGISEQ2000RS hardware using MGIEasy V3 chemistry for 100 bp paired-end sequencing at the Monash Micromon Genomics platform.

Raw Sequencing data was processed, and samples were demultiplexed using CellRanger multi (v.7.0) (17), with the CMOs provided (Supplemental Table 8). Reads were aligned to the Homo sapiens GRCh38 (2020-A) reference. Feature counts per cell, cell barcodes and mapping rate were used to evaluate cell quality. An average of 975 cells were capture per sample with high mapping rates to reference (>98%) and an average of 8,251 features per cell (Supplemental File 3). Quality control measures, preprocessing steps, and all subsequent analysis and related code is documented and available in a public github repository, which also features an interactive marker lookup site to interrogate gene expression within experimental groups and cell types https://github.com/MonashBioinformaticsPlatform/sc-hyp-org.

Further investigation of data quality and analysis was performed using R package Seurat (v. 4.3.0) (18). The distribution of the number of features per cell (unique UMIs per cell), number of reads per feature (UMI counts), and mitochondrial content was assessed to determine the quality of the cells. A threshold of cells with more than 200 features (unique UMIs per cell) was applied. Mitochondrial gene expression in kidney cells is higher than other organs (up to 50%) (19), and was expected to be increased in our experiment. Therefore, mitochondrial expression was not considered as sign of dying cells but a response of kidney cells to the conditions in this experiment and no mitochondrial percentage threshold was used to filter out cells. Normalization of reads per cells (UMI counts) was performed using the SCT transform method (20). An elbow plot of a Principal Component Analysis (PCA) was used to visualize dimensionality; 15 dimensions represented most of the variability. Dimensionality reduction was performed using Uniform Manifold Approximation and Projection (UMAP) with 15 dimensions.

The d20h group was profoundly affected by hypoxic exposure and formed unique clusters, while data from the d20n, d25n/n and d25 h/n groups clustered together. Various strategies were attempted to integrate the d20h group and enable comparison of cell-type specific changes in comparison to control data within the same cell type-cluster. A cluster tree (21) was generated to evaluate the clustering resolutions from 0 to 2. A resolution of 0.7 was selected as this resolution captured most of the clusters' variability without diverging into smaller subclusters with low number of cells (<18 cells) (Supplemental Figure 9A). Second, 30 to 50 of the most variable gene markers per cluster in the normoxic clustering were used as anchor genes to identify clusters in hypoxic groups. Finally, the cluster labelling was informed and refined by observing the segregation of the clusters, identification of gene markers in literature, comparison to the Enrichr database (22) and the use of the R package DevKidCC (23). Gene set enrichment analysis was used to identify clusters defined by signatures of proliferation and cellular stress, as well as known 'off-target' neural and myocyte-like populations (24). Changes in the proportion of cell populations was assessed using the propeller method from the R package speckle (25). Differentially Expressed Genes (DEGs) were identified using the pseudobulk approach (26). The pseudobulk analysis was performed using the R packages limma (27) and edgeR (28). ScRNAseq Sequencing data was submitted to GEO Dataset under the ID GSE236379.

**Preparation of Samples for Untargeted Metabolomic and Proteomic Analysis**

Without disrupting the organoid architecture, individual organoids were transferred to a low binding 1.5 mL Eppendorf tube. For each experimental group, eight replicate organoids were used for the metabolomic study and four replicates were used for the proteomic study. Organoids were washed with 1 mL of ice cold 0.9 % NaCl. After a 10 min centrifugation at 1000 x g, 4 °C, all the saline solution was removed and organoids frozen in liquid nitrogen. Frozen pellets were resuspended in 200 µL of extraction solvent consisting of 2:6:1 chloroform:methanol:water with a mixture of internal standards (CHAPS, CAPS, PIPES and TRIS; 2 µM each). Samples were frozen in liquid nitrogen and thawed on ice three times then agitated on a vibrating mixer for 20 minutes at 4°C before centrifugation at 20,000 × g and 4°C for 10 min. The supernatant was transferred to a glass vial for LC-MS based metabolomic analysis, while the pellet of four samples was used for proteomics analysis as described below.

Cell Lysis and Enzymatic Digestion

Protein pellets were resuspended in 1% (w/v) Sodium deoxycholate (SDC) (Sigma, D6750-100G), 100mM Tris pH 8.1 buffer, boiled at 95°C for 5 min. Subsequently, DNA was sheared using probe ultra-sonication at 50% amplitude (Q125 sonicator, Q-Sonica, Newtown, Connecticut, USA) with three rounds of 10-second intervals. Protein concentration was measured using a BCA kit according to the kit's manual (Pierce BCA Protein Assay Kit #2322). TCEP (500 mM Bond-Breaker Thermo, #77720) to a final concentration of 10 mM was added to denature samples and incubated for 30 min at 50°C. Then samples were alkylated by adding chloroacetamide to a concentration of 40 mM and incubate for 20 min at RT in the dark. Finally, 1:100 of trypsin (Promega, V5111) was added and incubated at 37°C with shaking overnight. Trypsin digestion was stopped by adding formic acid (Sigma, 56302-50ML-F) to a final concentration of 1%, and pH was ensured to be pH ~3. Samples were centrifuged at 5 min with max speed and supernatants were transferred to new tubes.

Stage tip SDB Clean-up Protocol

StageTiping was used to clean the peptide digest using SDB-RPS (Empore) (29). Samples were reconstituted in 20 $\mu$L of loading buffer (2% acetonitrile, 0.1% TFA and 97.9% MilliQ water) (Thermo Fisher Scientific, Optima LC/MS, A955-4) containing iRT peptides, sonicated for 10 minutes in a sonicator water bath, centrifuged at max speed for 5 minutes and transferred into MS vials to be loaded into the mass spectrometer.

Liquid Chromatography Mass Spectrometry for Proteomic Analysis

Using a Dionex UltiMate 3000 RSLCnano system equipped with a Dionex UltiMate 3000 RS autosampler, an Acclaim PepMap RSLC analytical column (75 µm x 50 cm, nanoViper, C18, 2 µm, 100Å; Thermo Scientific) and an Acclaim PepMap 100 trap column (100 µm x 2 cm, nanoViper, C18, 5 µm, 100Å; Thermo Scientific), the tryptic peptides were separated by increasing concentrations of 80% acetonitrile (ACN) / 0.1% formic acid at a flow of 250 nl/min for 158 min and analyzed with an Orbitrap Fusion Tribrid mass spectrometer (ThermoFisher Scientific). The instrument was operated in data dependent acquisition mode to automatically switch between full scan MS and MS/MS acquisition. Each survey full scan (m/z 375–1800) was acquired in the Orbitrap with a resolution of 120,000 (at m/z 200) after accumulating ions with a normalized AGC (automatic gain control) target of 250% and an automated maximum injection time. The cycle time was set to two seconds with the most intense multiply charged ions (z = 2-7) sequentially isolated and fragmented in the collision cell by higher-energy collisional dissociation (HCD) with a resolution of 30,000, an AGC target of 400% and an automated maximum injection time. Dynamic exclusion was set to 15 seconds.

Proteomic Mass Spectrometric Data Analysis

The mass spectrometric raw data files were analyzed with the Fragpipe software suite 17.1 (MSFragger version 3.4, Philosopher version 4.1.1 (30, 31) to obtain protein identifications and their respective label-free quantification (LFQ) values. The standard label free quantification match-between-runs (LFQ-MBR) workflow was applied with no changes to workflow, employing IonQuant (Version 1.7.17) (32) and the MaxLFQ method of protein abundance calculations (33). Searches were performed against the Human SwissProt proteome (accessed June 2020) and with common contaminants. The proteomics data were further analyzed using LFQ-Analyst which performed data manipulation and statistical tests with standard parameters (see (34)).

Liquid Chromatography Mass Spectrometry for Metabolomic Analysis

Untargeted metabolomic analysis was performed with a Dionex RSLC3000 UHPLC coupled to a Q-Exactive Plus MS (Thermo Fisher Scientific). The chromatography utilized a ZIC-p(HILIC) column 5µm, 150 x 4.6 mm (Merck Millipore, Australia) at 25 °C. A gradient elution of 20 mM ammonium carbonate (A) and acetonitrile (B) (linear gradient time-%B: 0 min-80%, 15 min-50%, 18 min-5%, 21 min-5%, 24 min-80%, 32 min-80%) was utilized. Flow rate was maintained at 300 µL/min. Samples were kept in the autosampler (6°C) and 10 µL was injected for analysis. MS was performed at 70,000 (at m/z = 200) resolution operating in rapid switching positive (4 kV) and negative (−3.5 kV) mode electrospray ionization (capillary temperature 300°C; sheath gas flow rate 50; auxiliary gas flow rate 20; sweep gas 2; probe temp 120°C). For accurate metabolite identification, a standard library of ~500 metabolites were analysed before sample testing and accurate retention time for each standard was recorded. This standard library also forms the basis of a retention time prediction model used to provide putative identification of metabolites not contained within the standard library (35). Acquired LC-MS data was processed in an untargeted fashion using open source software IDEOM, which initially used msConvert (ProteoWizard)(36) to convert raw LC-MS files to mzXML format and XCMS to pick peaks to convert to .peakML files (37). Mzmatch was subsequently used for sample alignment and filtering (38). On importing the aligned peak data into IDEOM, the intensity was normalised to the median peak intensity before running the identification macro to provide identification of the chromatographic features (37).

**Organoid Tissue Processing and Immunostaining**

Organoids were washed with DPBS (no calcium, no magnesium, Thermo Fisher Scientific). Forcibly pipetting DPBS around the organoids caused the organoids to detach from the transwell without causing tissue damage. Organoids were then placed in a low attachment 48-well plate (Thermo Fisher Scientific) with a cut P1000 tip. For whole mount staining, organoids were fixed in 2 % (paraformaldehyde) PFA for 20 min and blocked for 30 min (5 % Donkey serum, 0.1 % Triton X-100, made up in PBS) at room temperature. Organoids were then placed overnight with the primary antibody or staining reagent at 4 °C. Kidney organoids were stained with LTL-biotin (B-1325, dilution 1:500), GATA3 (Cell Sig – 5852, dilution 1:300), UMOD (8595-0054, dilution 1:300), MAFB (HPA005653, dilution 1:300), NPHS1 (AF4269-SP, dilution 1:300), PBX1 (Cell Sig – 4342, dilution 1:300), MEIS1 (ATM39795, dilution 1:300), PECAM1 (SAB4502167, dilution 1:300), KIM1 (AF1750, dilution 1:100), CK (cytokeratin, ab115959, dilution 1:300), ICAM1 (Cell Sig – 4915, dilution 1:300) and COL IV (1340-01, dilution 1:400). Organoids were then incubated overnight at 4 °C with the secondary antibody (all 1:1000 dilution): donkey anti-Rabbit IgG (Thermo Fisher Scientific, A1004200); donkey anti-Sheep IgG (Thermo Fisher Scientific, A21448); Alexa Fluor 568 streptavidin (Thermo Fisher Scientific, S11226); donkey anti-Mouse IgG (Thermo Fisher Scientific, A32787, A32766); donkey anti-Goat (Thermo Fisher Scientific, A32849). Fluorescence images were collected using a 3i CSU-W1 T2 Yokogawa Spinning disk confocal microscope. Images were pseudo-coloured to permit overlay, cropped, sized, and enhanced for contrast and brightness with Photoshop and Illustrator (Adobe Systems) or ImageJ (NIH). Images for ICAM1 and KIM1 were collected using the VS200 slide scanner (Olympus) and analysed using Imaris. A surface rendering was created for either cytokeratin positive structures or LTL positive structures and the Mean intensity of ICAM or KIM1 were calculated in these surfaces respectively. A one-way ANOVA with multiple comparison test was performed on Prism (GraphPad). Representative images were generated in FIJI with equal brightness and contrast modifications across all conditions for display purposes.

For paraffin embedded tissue, organoids were fixed in 2 % PFA at 4 °C overnight. Samples were processed using a Leica Peloris II Tissue Processor at Monash Histology Platform. Briefly, organoids were processed to paraffin wax through 80% Ethanol and multiple stations of 100% Ethanol. Tissues were then dehydrated in several changes of Xylene before being infiltrated through multiple stations of Paraplast paraffin wax and embedded in the same. Immunostaining for collagen type I (Cell Sig – 72026, dilution 1:200) and PAX8 (MA5-32382, dilution 1:400) was performed on 4 µm thick sections using antigen retrieval with 0.1 mol/L sodium citrate, pH 6.0, and a three-layer avidin-biotin peroxidase complex staining method (Vector Laboratories), as described (39). Tubular damage was assessed on organoids using Periodic acid-Schiff (PAS) stain in sections (2 mm thick) of PFA-fixed tissue. Optimal cutting temperature compound (OCT)-embedded frozen sections was used for Oil Red O staining (40). Images were enhanced for contrast and brightness with Photoshop and Illustrator (Adobe Systems).

**Spatial Transcriptomics**

Kidney organoids +/- integrated iMacs were generated and subjected to hypoxic injury as described above. Organoids were embedded in HistoGel (Epredia, cat. no. HG-4000-012) as per the

Manufacturer's instructions, then processed for paraffin embedding (Ethanol 1 min x5, Xylene 1 min x3, Paraffin x3 17 mins). 5mm paraffin sections were cut from 2-5 organoids per experimental group and mounted directly onto a Xenium slide. Samples were profiled with the 380-gene Xenium Human Immuno-Oncology panel, using the Xenium Cell Segmentation kit (10X Genomics) at the Walter and Eliza Hall Institute Spatial Omics Platform Laboratory, as per the manufacturer's protocols. Raw counts from the 10x Xenium Onboard Analysis software were imported into R, loaded into a Seurat object, and normalized with SCTransform (v. 4.3.0) (37). Principal Component Analysis was performed on the SCT-normalized data, and the top 100 components were used for UMAP visualization. Cell type identities were assigned to Xenium clusters by integrating our organoid scRNAseq reference and spatial data using RCTD (41) and Seurat's anchor-based transfer, with results cross-validated and refined by manual assessment of clusters and marker gene expression on tissue images. Unsupervised graph-based clustering was performed in Seurat at multiple resolutions; final assignments using $\rho = 0.8$ clusters for nephron cells and $\rho = 0.3$ for other cell types. Pseudobulk differential expression (DE) analysis was performed by aggregating UMI counts across cells per replicate. Pseudo-bulk counts were $\log_2$-transformed, quantile-normalized, and precision-weighted with voom (limma) (27, 42). Linear models with empirical Bayes moderation (43) were used to identify DE genes at FDR $\leq 0.05$ for comparisons at the level of sample, cell lineage. Cluster-based DE was performed on the single cell level with an FDR cutoff for DE $\leq 0.2$. Targeted DE analysis was performed on twenty-four manually defined Regions of Interest (ROIs) containing injured nephrons +/-iMacs in the d25h/n_iMacs organoid group, with cells selected by centroid location. The same pseudo-bulk limma-voom pipeline (FDR $\leq 0.05$) was used to compare all cells, non-iMac cells, and nephron cells only between +/-iMac ROIs. ROIs with fewer than three relevant cells were excluded.

### Supplemental Figures

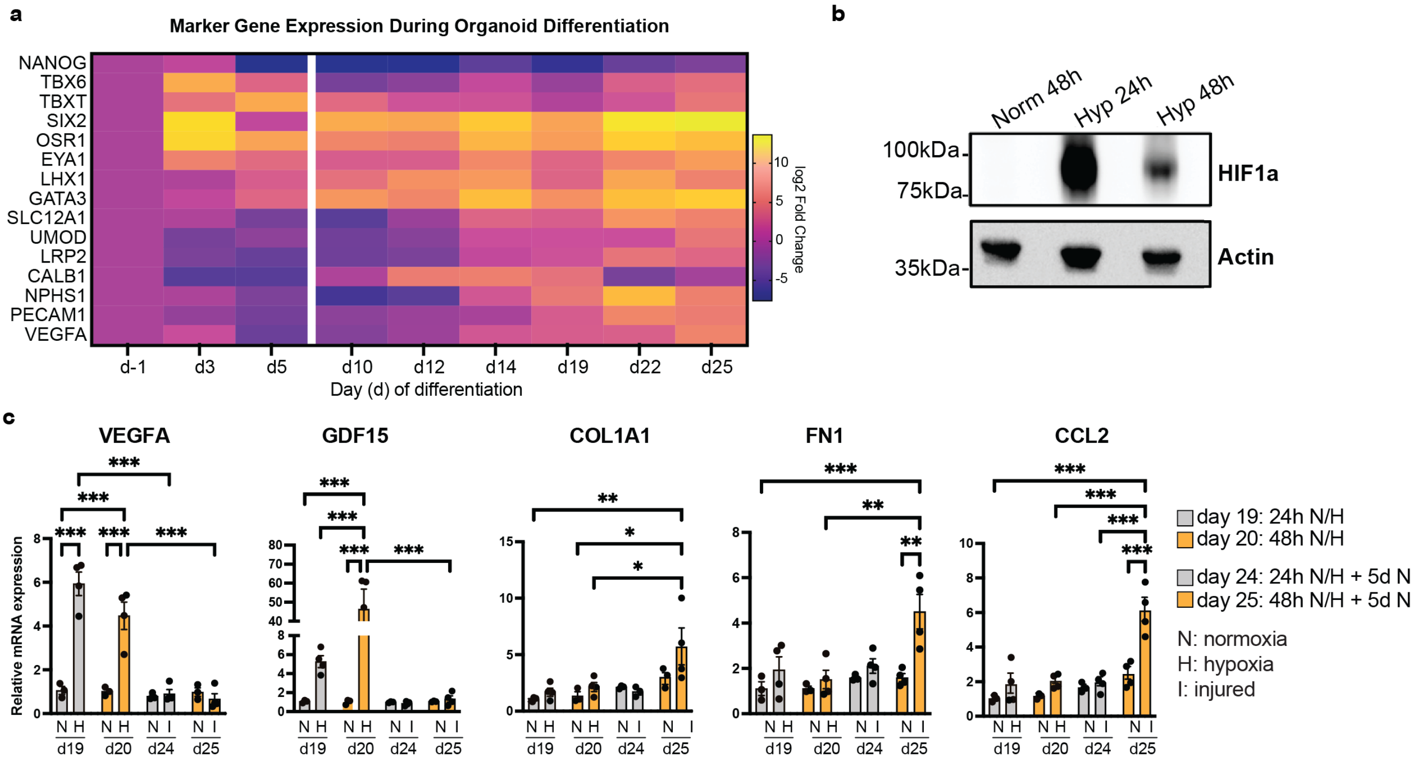


**Supplemental Figure 1. Kidney organoid development and hypoxic injury**. (**a**) Analysis of gene expression in kidney organoids from day -1 to day 25. Three wells from a six well-plate were used to determine gene expression on days 3 and 5. From day 10 to day 25, four organoids were used to determine gene expression levels. Data is represented as log2 fold change and normalised to *ACTB*. (**b**) Western blot results of HIF1A protein stabilization in nuclear extracts from organoids exposed to 24h and 48h of hypoxia compared to normoxic controls. β-actin was used as the loading control. (**b**) Comparison of the hypoxic injury response in kidney organoids subjected to 24 h (harvested at d19) and 48 h of hypoxia (harvested at d20), followed by a 5-day recovery period for both groups. qPCR results are shown as mean with SD; n=3-4. Y axis shows mRNA expression normalised to *ACTB* within each sample, relative to d19n mean. Analyses were performed by one-way ANOVA with Tukey's multiple comparison test: $*p < 0.05$, $**p < 0.01$, $***p < 0.001$.

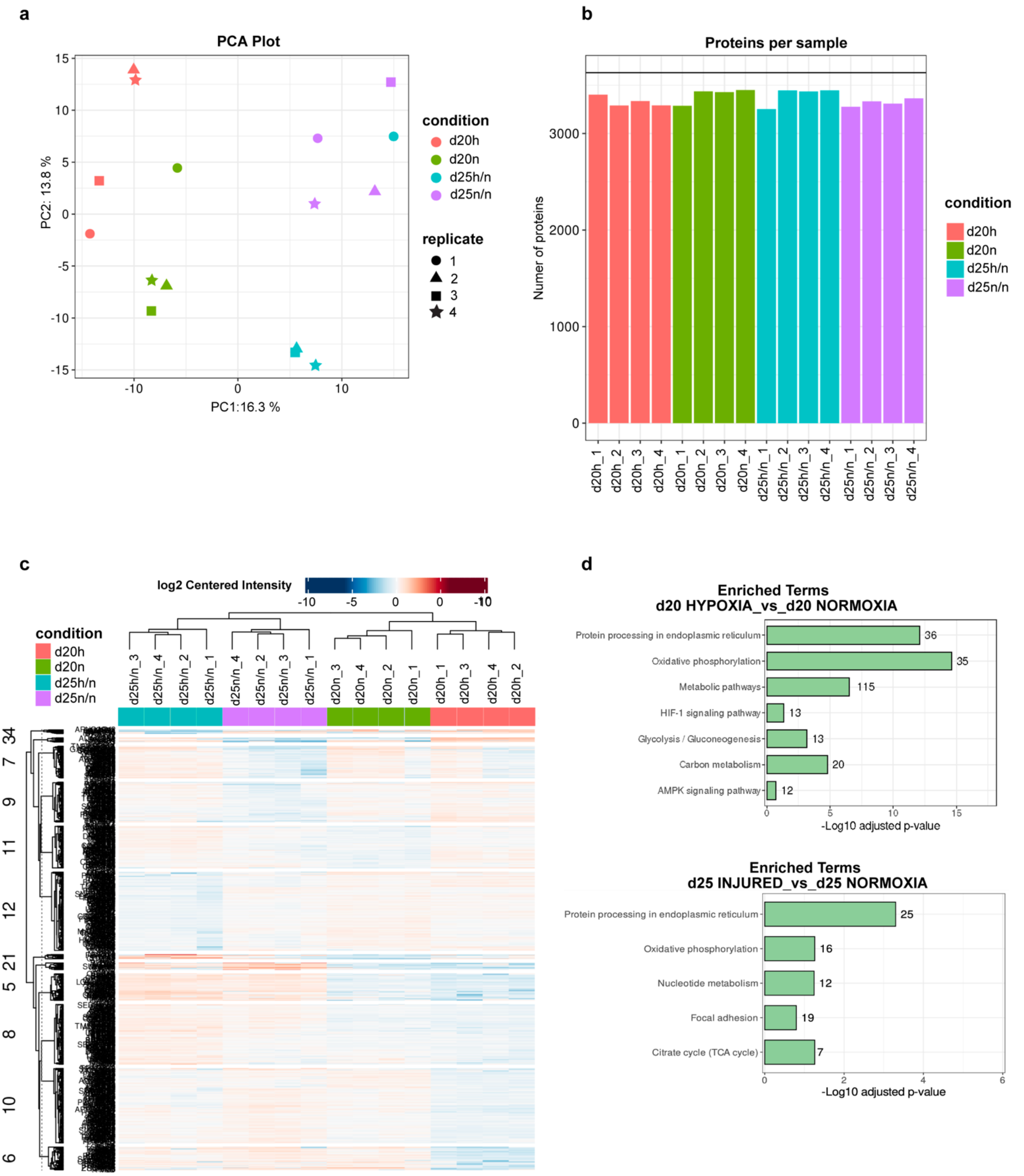


**Supplemental Figure 2. Proteomic analysis.** (**a**) PCA of the four replicates per condition. (**b**) Protein quantification per sample, after pre-processing. (**c**) Differential expression analysis in the 12 clusters. Heatmap repressing an overview of expression of all significant proteins (rows) in all samples (columns). (**d**) KEGG pathway enrichment analysis using the web-based DAVID software (p-value < 0.05) for differentially expressed genes (BH adjusted p-value < 0.05 and absolute LogFC > 0.05). *P*-values were corrected with Benjamini Hochberg approach.

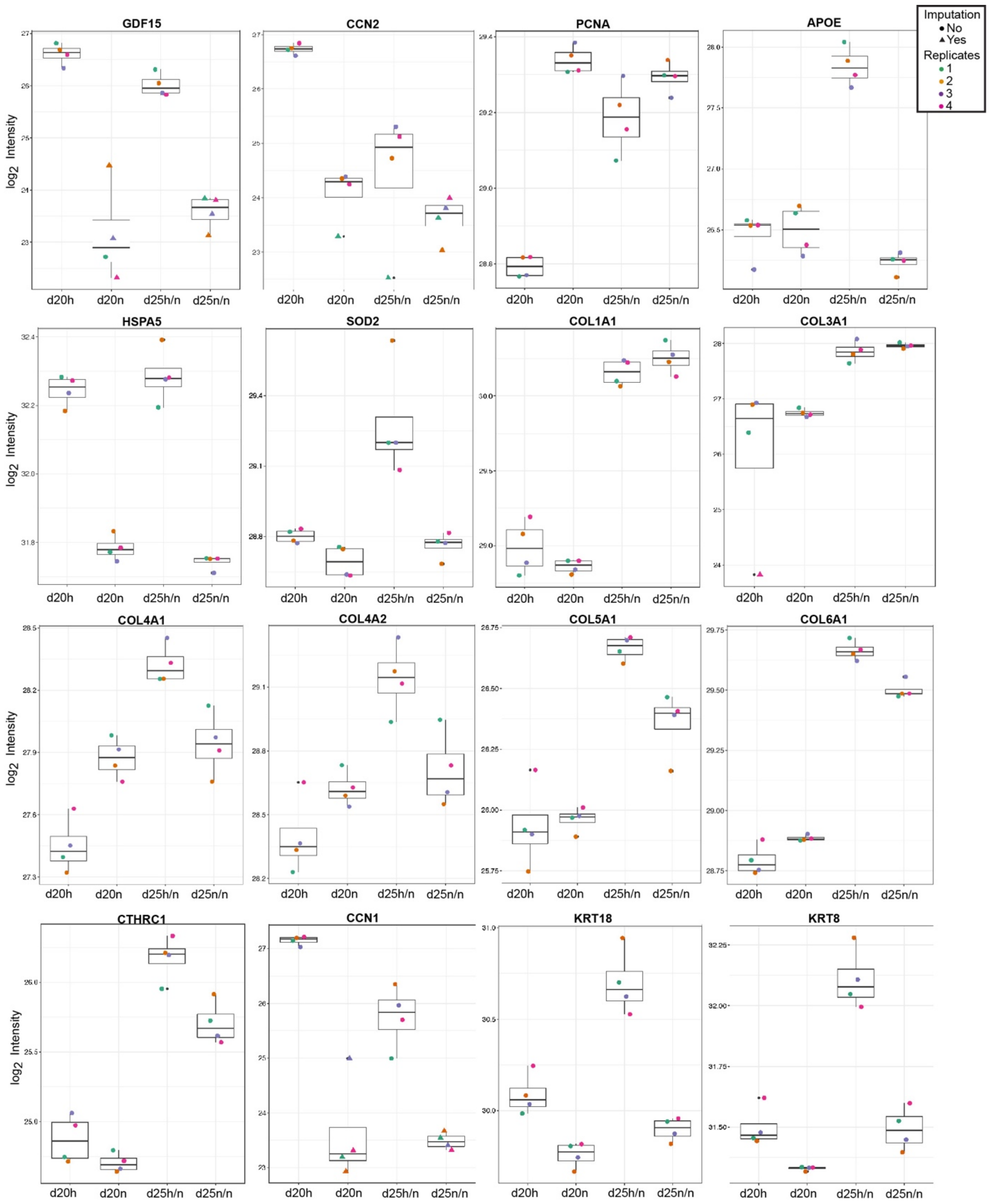


**Supplemental Figure 3. Key changes in proteins associated with AKI and maladaptive regeneration.** Data produced using LFQ-Analyst Proteins. Illustrated proteins associated to kidney injury (GDF15, CCN1, KRT18, KRT8), hypoxia (CCN2, SOD2), cell cycle (PCNA), inflammation (APOE), UPR response (HSPA5), and ECM remodelling (COL1A1, COL3A1, COL4A1, COL4A2, COL5A1, COL6A1, CTHRC1).

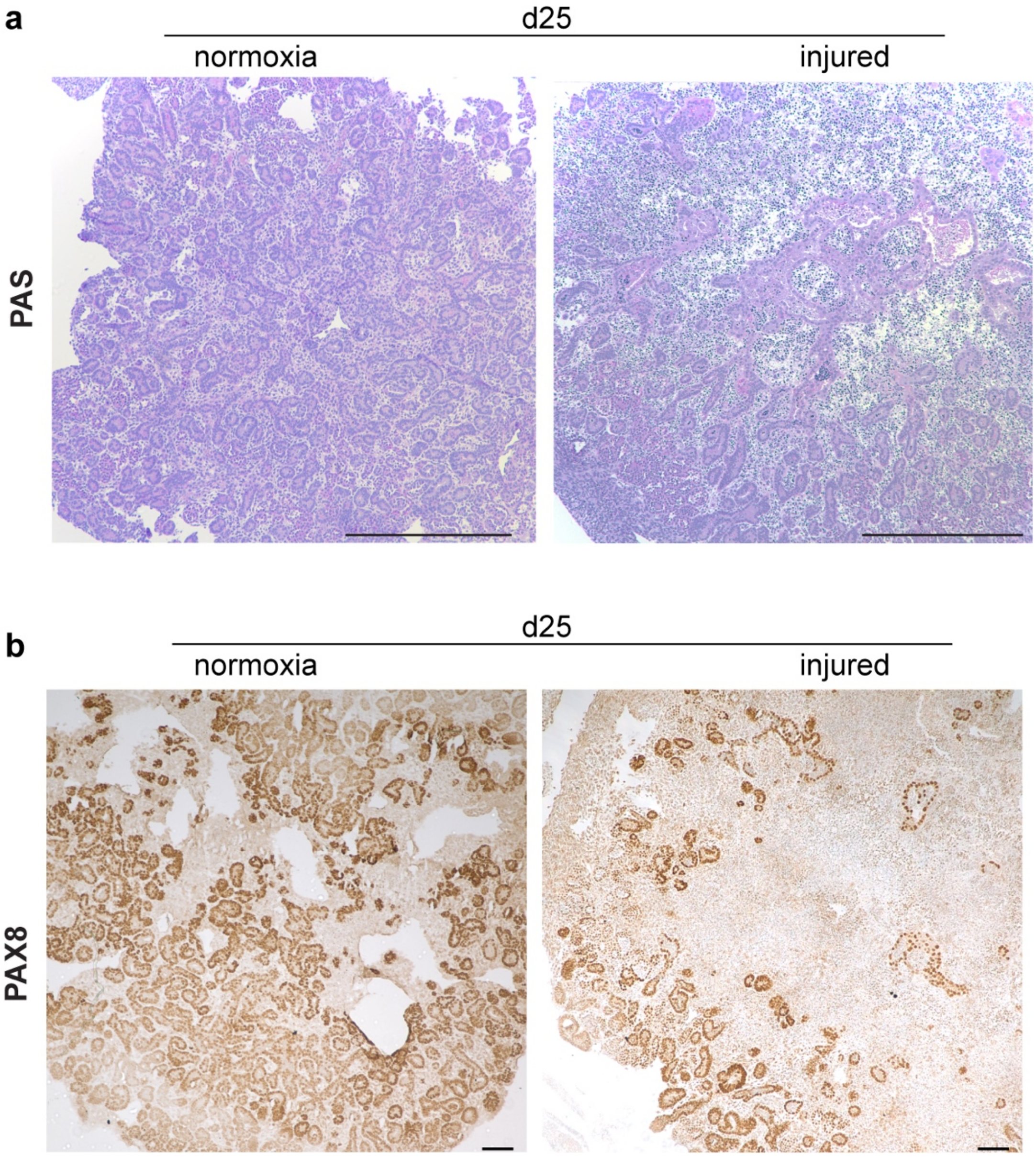


**Supplemental Figure 4. Immunostaining on sections of d25n/n and d25h/n organoids.** (**a**) Representative image (n=4) of Periodic acid-Schiff (PAS) staining showing a reduction in tubular structures in d25 injured organoids compared to controls. Scale bar = 500 µm. (**b**) Representative image (n=4) of PAX8 staining (epithelial marker) showing reduction tubular structures in injured organoids. Scale bar = 100 µm.

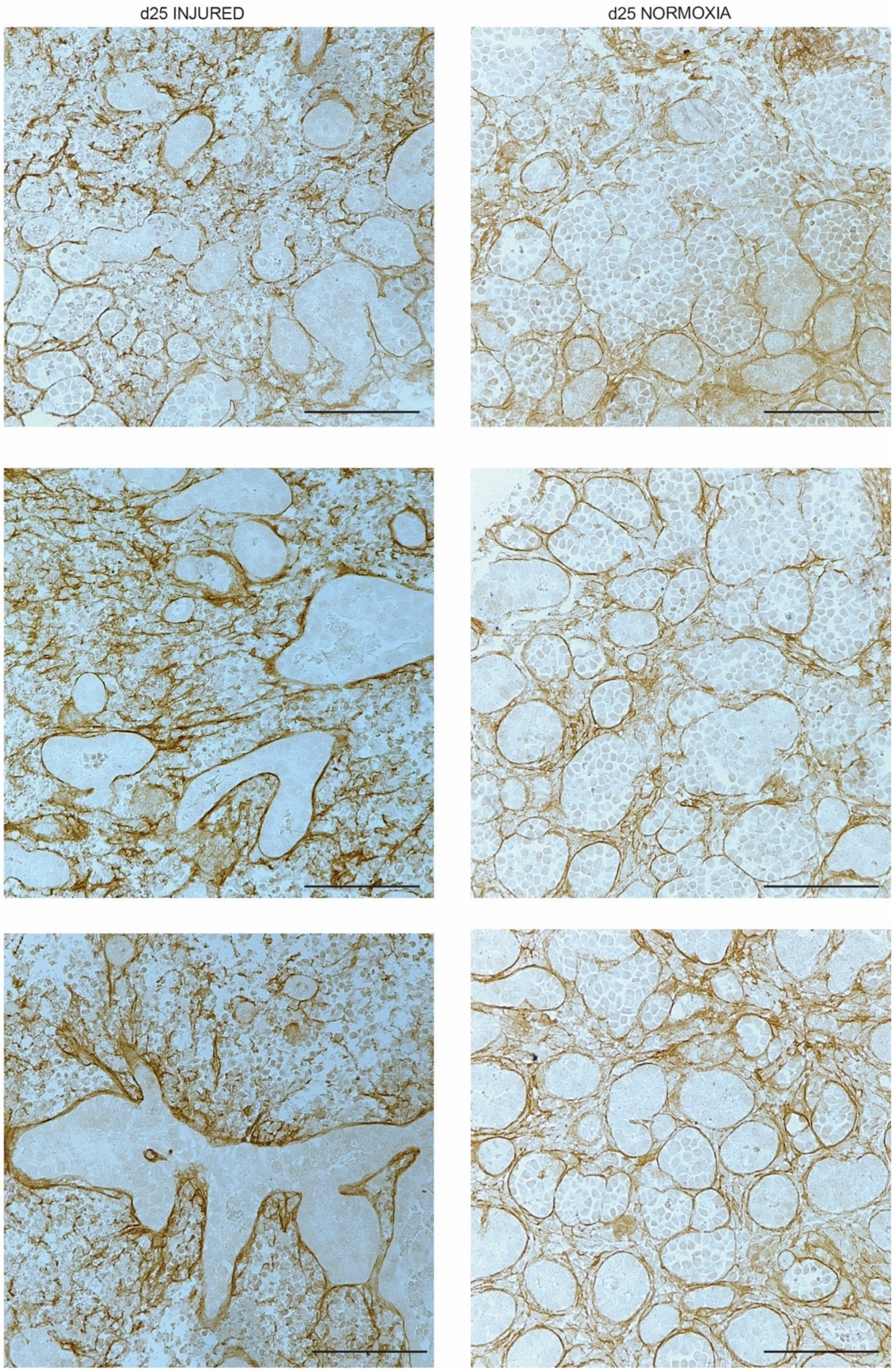


**Supplemental Figure 5. Collagen type I staining on sections of d25 injured and d25 control organoids.** Staining shows disruption of tubular morphology in three separate organoids. Scale bars = 500 µm.

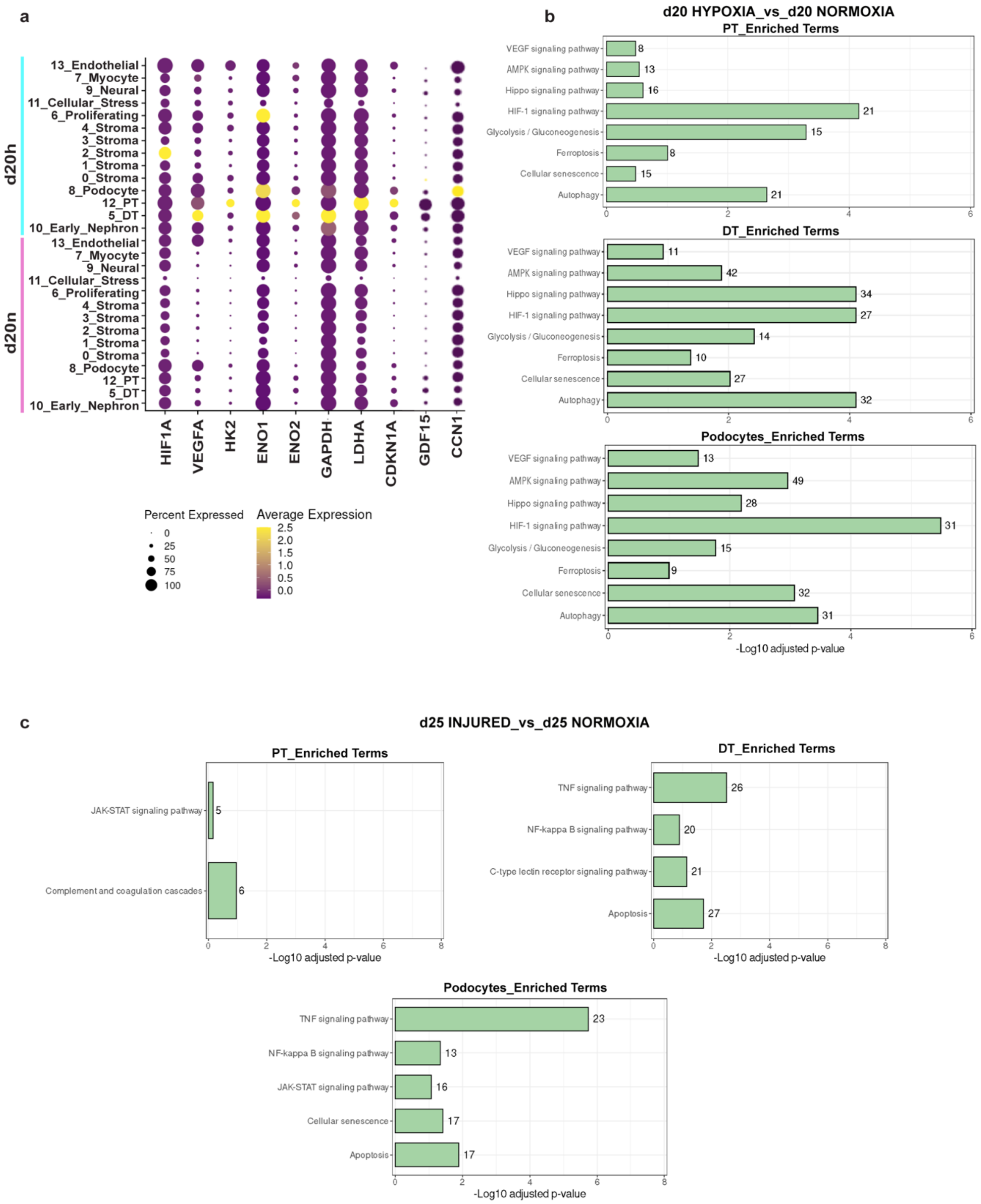


**Supplemental Figure 6. Hypoxia induces profound transcriptional changes in PT and DT of kidney organoids.** (**a**) Dot plot of the expression of hypoxia-specific genes in the 14 clusters of hypoxic and normoxic d20 organoids. (**b**) Commonly enriched KEGG pathways in PT (proximal tubule), DT (distal tubule) and podocytes of hypoxic d20 organoids compared to controls. (**c**) Commonly enriched KEGG pathways in PT, DT and podocytes of d25 injured organoids compared to controls. Pathways obtained using the web-based DAVID software (adjusted p-value < 0.05) for differentially expressed genes (BH adjusted p-value < 0.05 and absolute LogFC > 0.05). *P*-values were corrected with Benjamini Hochberg approach.

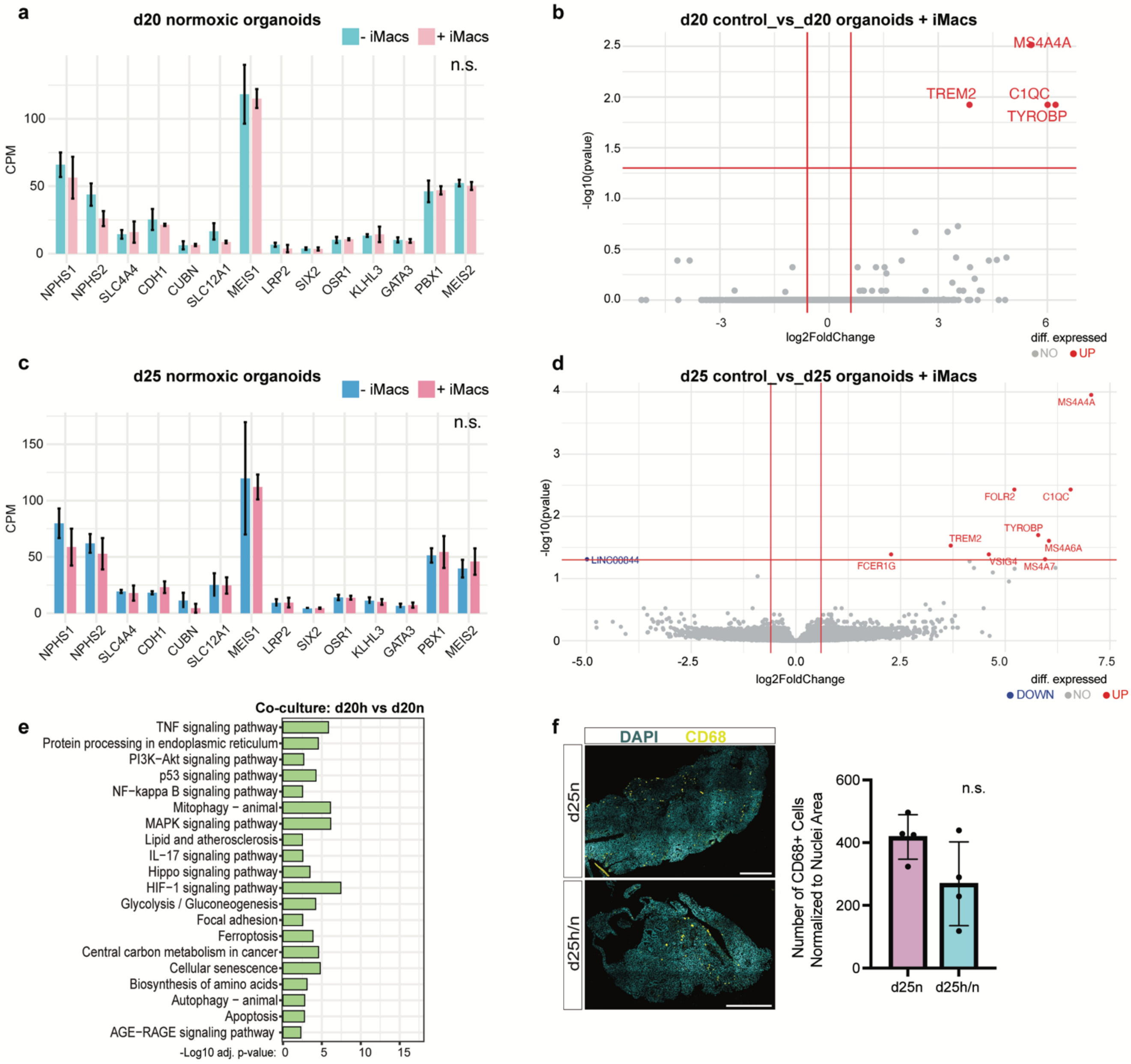


**Supplemental Figure 7. Bulk RNAseq analysis of kidney organoids containing iMacs.** (**a**) Expression levels of cell type markers in d20 normoxic organoids with- and without-macrophages (+/-iMacs). Error bars represent standard deviation of 3 biological replicates. Two-tailed t-test was performed, n = 3; ns. non-significant. (**b**) Volcano plot of DE genes (LogFC>1, adj. p<0.05) between d20 normoxic organoids +/-iMacs. Red indicates upregulated genes in +iMac group; non-significant changes in grey. (**c**) Expression levels of cell type markers in d25 normoxic organoids +/-iMacs (n=3). (**d**) DE genes between d25 normoxic organoids with and without macrophages (LogFC>1, adj. p<0.05). (**e**) Upregulated pathways in d20 iMac-organoids after hypoxia. DE genes (p < 0.05) were used to perform enrichment analysis with DAVID; pathways with FDR<0.05 displayed. (**f**) Representative images of d25 hypoxic and normoxic organoids +iMacs labelled with CD68. Scale bar = 500 µm. Quantification of CD68 positive cells normalized to DAPI, n=4, ns. non-significant.

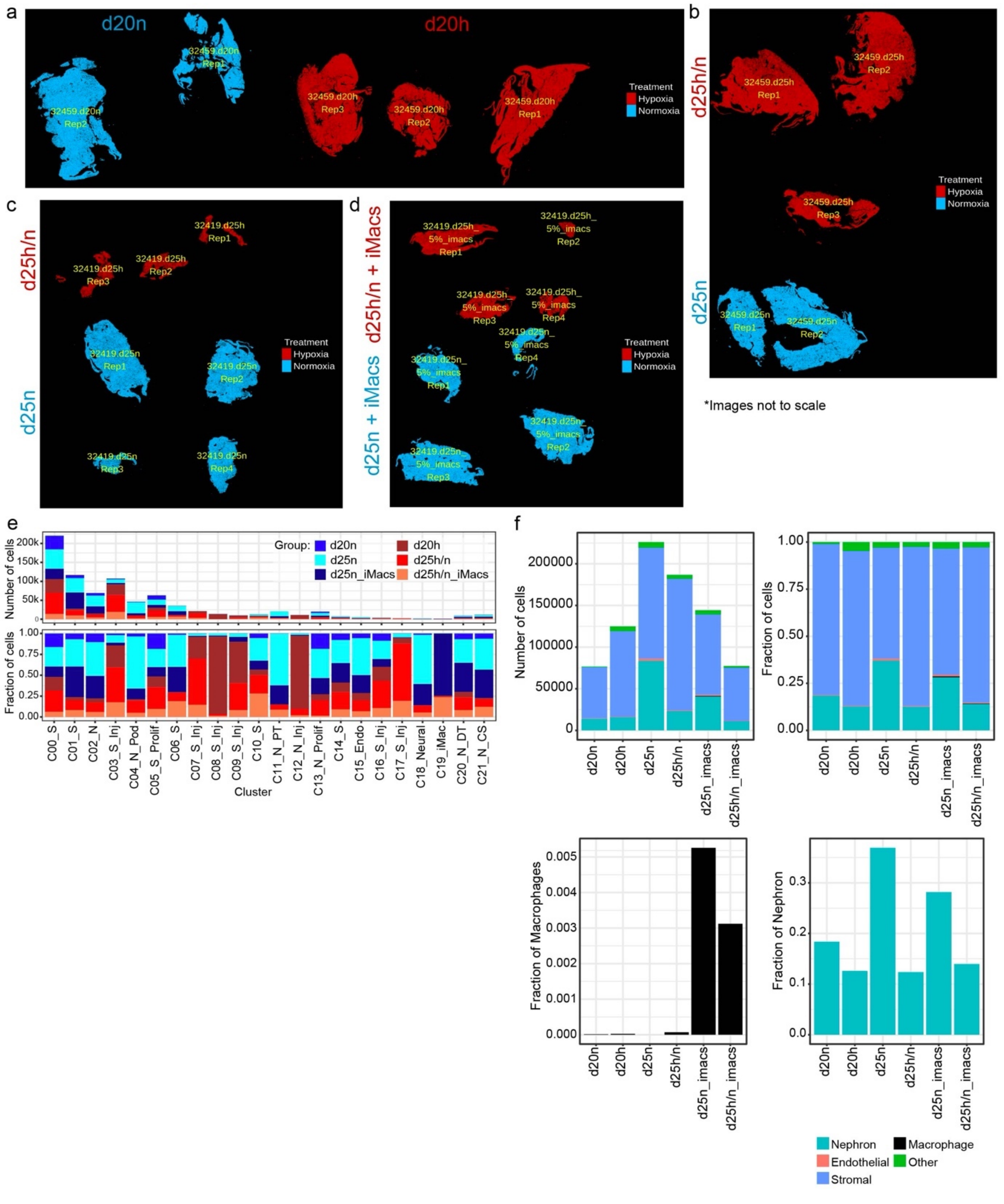

**Supplemental Figure 8. Xenium spatial analysis of kidney organoids co-cultured with iMacs.** (**a-d**) Images of tissue outlines and experimental groups for all spatial data. Stage and experimental groups indicated as per labels, images not to scale. (**e**) Absolute number of cells and fractional contributions to each cluster from each sample group according to colours and labels in the colour key. (**f**) Number and fraction of cells in each major cell type or lineage by sample group.

**Supplemental Table 1.** Cell Proportions: hypoxia_d20 vs control_d20 (t-test)

| BaselineProp.clusters | BaselineProp.Freq | PropMean D20h | PropMean D20n | P.Value | FDR |
|---|---|---|---|---|---|
| 13_Endothelial | 0.0252 | 0.0038 | 0.0649 | 0.0000 | 0.0000 |
| 11_Cellular_Stress | 0.0423 | 0.0531 | 0.0229 | 0.0001 | 0.0005 |
| 0_Stroma | 0.0947 | 0.1154 | 0.0566 | 0.0003 | 0.0012 |
| 8_Podocyte | 0.0617 | 0.0740 | 0.0405 | 0.0032 | 0.0114 |
| 5_Distal_Tubule | 0.1234 | 0.1420 | 0.0887 | 0.0082 | 0.0205 |
| 4_Stroma | 0.0973 | 0.1121 | 0.0695 | 0.0096 | 0.0205 |
| 6_Proliferating | 0.0695 | 0.0571 | 0.0930 | 0.0103 | 0.0205 |
| 1_Stroma | 0.1031 | 0.0899 | 0.1269 | 0.0431 | 0.0737 |
| 3_Stroma | 0.0864 | 0.0751 | 0.1071 | 0.0474 | 0.0737 |
| 2_Stroma | 0.1123 | 0.0987 | 0.1365 | 0.0635 | 0.0889 |
| 9_Neural | 0.0423 | 0.0379 | 0.0504 | 0.1409 | 0.1794 |
| 12_Proximal_Tubule | 0.0229 | 0.0267 | 0.0165 | 0.2808 | 0.3276 |
| 7_Myocyte | 0.0595 | 0.0567 | 0.0637 | 0.4149 | 0.4469 |
| 10_Early_Nephron | 0.0593 | 0.0574 | 0.0627 | 0.6153 | 0.6153 |

**Supplemental Table 2.** Cell Proportions: injured_d25 vs control_d25 (t-test)

| BaselineProp.clusters | BaselineProp.Freq | PropMean D25h/n | PropMean D25n/n | P.Value | FDR |
|---|---|---|---|---|---|
| 10_Early_Nephron | 0.0289 | 0.0187 | 0.0547 | 0.0001 | 0.0008 |
| 0_Stroma | 0.1549 | 0.1827 | 0.0830 | 0.0004 | 0.0025 |
| 4_Stroma | 0.0889 | 0.0675 | 0.1429 | 0.0008 | 0.0038 |
| 9_Neural | 0.0539 | 0.0400 | 0.0883 | 0.0026 | 0.0091 |
| 11_Cellular_Stress | 0.0389 | 0.0459 | 0.0195 | 0.0033 | 0.0092 |
| 13_Endothelial | 0.0134 | 0.0089 | 0.0239 | 0.0052 | 0.0122 |
| 5_Distal_Tubule | 0.0598 | 0.0495 | 0.0851 | 0.0154 | 0.0309 |
| 6_Proliferating | 0.0949 | 0.1059 | 0.0668 | 0.0198 | 0.0324 |
| 12_Proximal_Tubule | 0.0136 | 0.0098 | 0.0236 | 0.0208 | 0.0324 |
| 1_Stroma | 0.1317 | 0.1471 | 0.0946 | 0.0289 | 0.0405 |
| 8_Podocyte | 0.0337 | 0.0360 | 0.0288 | 0.2566 | 0.3209 |
| 7_Myocyte | 0.0839 | 0.0883 | 0.0726 | 0.2751 | 0.3209 |
| 2_Stroma | 0.0990 | 0.0958 | 0.1091 | 0.4671 | 0.5030 |
| 3_Stroma | 0.1044 | 0.1039 | 0.1073 | 0.9145 | 0.9145 |

#### Supplemental References


1. Howden SE, Little MH. Generating Kidney Organoids from Human Pluripotent Stem Cells Using Defined Conditions. Methods Mol Biol. 2020;2155:183-92.
2. Takasato M, Er PX, Becroft M, Vanslambrouck JM, Stanley EG, Elefanty AG, et al. Directing human embryonic stem cell differentiation towards a renal lineage generates a self-organizing kidney. Nature Cell Biology. 2014;16(1):118-26.
3. Takasato M, Er PX, Chiu HS, Maier B, Baillie GJ, Ferguson C, et al. Kidney organoids from human iPS cells contain multiple lineages and model human nephrogenesis. Nature. 2015;526(7574):564-8.
4. Berrocal-Rubio MA, Pawer YDJ, Dinevska M, De Paoli-Iseppi R, Widodo SS, Gleeson J, et al. Discovery of NRG1-VII: the myeloid-derived class of NRG1. BMC Genomics. 2024;25(1):814.
5. Rajab N, Angel PW, Deng Y, Gu J, Jameson V, Kurowska-Stolarska M, et al. An integrated analysis of human myeloid cells identifies gaps in in vitro models of in vivo biology. Stem Cell Reports. 2021;16(6):1629-43.
6. Dobin A, Davis CA, Schlesinger F, Drenkow J, Zaleski C, Jha S, et al. STAR: ultrafast universal RNA-seq aligner. Bioinformatics. 2013;29(1):15-21.
7. Li H, Handsaker B, Wysoker A, Fennell T, Ruan J, Homer N, et al. The sequence alignment/map format and SAMtools. bioinformatics. 2009;25(16):2078-9.
8. Smith T, Heger A, Sudbery I. UMI-tools: modeling sequencing errors in Unique Molecular Identifiers to improve quantification accuracy. Genome research. 2017;27(3):491-9.
9. Liao Y, Smyth GK, Shi W. featureCounts: an efficient general purpose program for assigning sequence reads to genomic features. Bioinformatics. 2014;30(7):923-30.
10. Durinck S, Spellman PT, Birney E, Huber W. Mapping identifiers for the integration of genomic datasets with the R/Bioconductor package biomaRt. Nature protocols. 2009;4(8):1184-91.
11. Ritchie ME, Phipson B, Wu D, Hu Y, Law CW, Shi W, et al. limma powers differential expression analyses for RNA-sequencing and microarray studies. Nucleic acids research. 2015;43(7):e47-e.
12. Law CW, Chen Y, Shi W, Smyth GK. voom: Precision weights unlock linear model analysis tools for RNA-seq read counts. Genome biology. 2014;15(2):1-17.
13. Powell D. Degust: interactive RNA-seq analysis. Drpowell/Degust. 2015;4(1):4.1.
14. Sherman BT, Hao M, Qiu J, Jiao X, Baseler MW, Lane HC, et al. DAVID: a web server for functional enrichment analysis and functional annotation of gene lists (2021 update). Nucleic acids research. 2022;50(W1):W216-W21.
15. Subramanian A, Tamayo P, Mootha VK, Mukherjee S, Ebert BL, Gillette MA, et al. Gene set enrichment analysis: a knowledge-based approach for interpreting genome-wide expression profiles. Proceedings of the National Academy of Sciences. 2005;102(43):15545-50.
16. Liberzon A, Subramanian A, Pinchback R, Thorvaldsdóttir H, Tamayo P, Mesirov JP. Molecular signatures database (MSigDB) 3.0. Bioinformatics. 2011;27(12):1739-40.
17. Zheng GX, Terry JM, Belgrader P, Ryvkin P, Bent ZW, Wilson R, et al. Massively parallel digital transcriptional profiling of single cells. Nat Commun. 2017;8:14049.
18. Hao Y, Hao S, Andersen-Nissen E, Mauck WM, 3rd, Zheng S, Butler A, et al. Integrated analysis of multimodal single-cell data. Cell. 2021;184(13):3573-87 e29.
19. Liao J, Yu Z, Chen Y, Bao M, Zou C, Zhang H, et al. Single-cell RNA sequencing of human kidney. Sci Data. 2020;7(1):4.
20. Hafemeister C, Satija R. Normalization and variance stabilization of single-cell RNA-seq data using regularized negative binomial regression. Genome Biol. 2019;20(1):296.
21. Zappia L, Oshlack A. Clustering trees: a visualization for evaluating clusterings at multiple resolutions. Gigascience. 2018;7(7).
22. Xie Z, Bailey A, Kuleshov MV, Clarke DJB, Evangelista JE, Jenkins SL, et al. Gene Set Knowledge Discovery with Enrichr. Current Protocols. 2021;1(3):e90.

23. Wilson SB, Howden SE, Vanslambrouck JM, Dorison A, Alquicira-Hernandez J, Powell JE, et al. DevKidCC allows for robust classification and direct comparisons of kidney organoid datasets. Genome Med. 2022;14(1):19.
24. Xie Z, Bailey A, Kuleshov MV, Clarke DJB, Evangelista JE, Jenkins SL, et al. Gene Set Knowledge Discovery with Enrichr. Curr Protoc. 2021;1(3):e90.
25. Phipson B, Sim CB, Porrello ER, Hewitt AW, Powell J, Oshlack A. propeller: testing for differences in cell type proportions in single cell data. Bioinformatics. 2022;38(20):4720-6.
26. Squair JW, Gautier M, Kathe C, Anderson MA, James ND, Hutson TH, et al. Confronting false discoveries in single-cell differential expression. Nat Commun. 2021;12(1):5692.
27. Ritchie ME, Phipson B, Wu D, Hu Y, Law CW, Shi W, et al. limma powers differential expression analyses for RNA-sequencing and microarray studies. Nucleic Acids Res. 2015;43(7):e47.
28. Robinson MD, McCarthy DJ, Smyth GK. edgeR: a Bioconductor package for differential expression analysis of digital gene expression data. Bioinformatics. 2010;26(1):139-40.
29. Humphrey SJ, Karayel O, James DE, Mann M. High-throughput and high-sensitivity phosphoproteomics with the EasyPhos platform. Nature Protocols. 2018;13(9):1897-916.
30. Teo GC, Polasky DA, Yu F, Nesvizhskii AI. Fast Deisotoping Algorithm and Its Implementation in the MSFragger Search Engine. J Proteome Res. 2021;20(1):498-505.
31. Kong AT, Leprevost FV, Avtonomov DM, Mellacheruvu D, Nesvizhskii AI. MSFragger: ultrafast and comprehensive peptide identification in mass spectrometry–based proteomics. Nature Methods. 2017;14(5):513-20.
32. Yu F, Haynes SE, Nesvizhskii AI. IonQuant Enables Accurate and Sensitive Label-Free Quantification With FDR-Controlled Match-Between-Runs. Mol Cell Proteomics. 2021;20:100077.
33. Cox J, Neuhauser N, Michalski A, Scheltema RA, Olsen JV, Mann M. Andromeda: a peptide search engine integrated into the MaxQuant environment. J Proteome Res. 2011;10(4):1794-805.
34. Shah AD, Goode RJA, Huang C, Powell DR, Schittenhelm RB. LFQ-Analyst: An Easy-To-Use Interactive Web Platform To Analyze and Visualize Label-Free Proteomics Data Preprocessed with MaxQuant. Journal of Proteome Research. 2020;19(1):204-11.
35. Creek DJ, Jankevics A, Breitling R, Watson DG, Barrett MP, Burgess KEV. Toward Global Metabolomics Analysis with Hydrophilic Interaction Liquid Chromatography–Mass Spectrometry: Improved Metabolite Identification by Retention Time Prediction. Analytical Chemistry. 2011;83(22):8703-10.
36. Chambers MC, Maclean B, Burke R, Amodei D, Ruderman DL, Neumann S, et al. A cross-platform toolkit for mass spectrometry and proteomics. Nature Biotechnology. 2012;30(10):918-20.
37. Creek DJ, Jankevics A, Burgess KE, Breitling R, Barrett MP. IDEOM: an Excel interface for analysis of LC-MS-based metabolomics data. Bioinformatics. 2012;28(7):1048-9.
38. Scheltema RA, Jankevics A, Jansen RC, Swertz MA, Breitling R. PeakML/mzMatch: a file format, Java library, R library, and tool-chain for mass spectrometry data analysis. Anal Chem. 2011;83(7):2786-93.
39. Masaki T, Foti R, Hill PA, Ikezumi Y, Atkins RC, Nikolic-Paterson DJ. Activation of the ERK pathway precedes tubular proliferation in the obstructed rat kidney. Kidney Int. 2003;63(4):1256-64.
40. Cowan CS, Renner M, De Gennaro M, Gross-Scherf B, Goldblum D, Hou Y, et al. Cell Types of the Human Retina and Its Organoids at Single-Cell Resolution. Cell. 2020;182(6):1623-40.e34.
41. Cable DM, Murray E, Zou LS, Goeva A, Macosko EZ, Chen F, et al. Robust decomposition of cell type mixtures in spatial transcriptomics. Nat Biotechnol. 2022;40(4):517-26.
42. Law CW, Chen Y, Shi W, Smyth GK. voom: Precision weights unlock linear model analysis tools for RNA-seq read counts. Genome Biol. 2014;15(2):R29.
43. Smyth GK. Linear models and empirical bayes methods for assessing differential expression in microarray experiments. Stat Appl Genet Mol Biol. 2004;3:Article3.
